# Pervasive noise in human splice site selection

**DOI:** 10.1101/2025.07.16.665169

**Authors:** Eraj S. Khokhar, Kaitlyn Brokaw, Zachary J. Kartje, Valeria Sanabria, Nida Javeed, Ayush Kumar, Jonathan K. Watts, Athma A. Pai

**Affiliations:** RNA Therapeutics Institute, University of Massachusetts Chan Medical School, Worcester, MA; Department of Molecular, Cellular, and Cancer Biology, University of Massachusetts Chan Medical School, Worcester, MA

**Author notes:** These authors contributed equally (Eraj S. Khokhar, Kaitlyn Brokaw). Alexion Pharmaceuticals (AstraZeneca Rare Disease), Cambridge, MA.

## Abstract

RNA splicing has historically been thought to be highly efficient and accurate, with little opportunity for deviation from regulated alternative splicing decisions. This dogma has been challenged by recent observations that suggest that biological noise may contribute substantially to transcriptome diversity. However, quantitative understanding of stochastic variations in splicing is challenging because these transcripts are likely subject to rapid degradation. Here, we use ultra-deep sequencing across RNA compartments to track splicing intermediates in human cells and see abundant cryptic splicing associated with genomic features that promote splicing noise. We observe pervasive usage of low-fidelity splice sites, likely due to stochasticity in recruitment or binding of the spliceosome. These sites are most likely degraded in the nucleus rather than targeted by translation-dependent degradation processes, suggesting widespread surveillance and rapid quality control of non-productive RNA transcripts. Our findings provide unprecedented insights into the propensity for error in RNA processing mechanisms and the regulation of alternative splice sites across a gene.

## INTRODUCTION

Messenger RNA (mRNA) splicing is a crucial step in metazoan gene regulation, with 95% of genes containing introns that are excised to produce a mature transcript. In mammals, more than 90% of genes are alternatively spliced, most in a cell type or condition specific manner. Intron excision is performed by the spliceosome, a highly conserved molecular machine assembled from numerous small nuclear RNAs (snRNAs) and proteins. Spliceosome recruitment to pre-mRNA molecules is initiated by binding of U1 snRNA to the 5′ splicing donor site (GT or GC), U2 sRNA to the branchpoint A, and U2AF1 to the 3′ splicing acceptor site (AG or AC), with a total of at least ∼30 required nucleotides that are often separated by thousands of nucleotides. Less than 1% of mammalian introns are spliced using the minor spliceosome, which is regulated by U11 snRNA binding to a AT 5′ splicing donor, U12 snRNA to the branchpoint A, and ZRSR2 to the AC 3′ splicing acceptor site. Both major and minor spliceosome regulated splice sites are among the most conserved metazoan genetic elements,^1,2^ leading to a prevalent perspective that the spliceosome is a high-fidelity, deterministic enzyme.^3^ However, core dinucleotides occur within longer, more degenerate consensus motifs that contribute to binding strength.^4^ Recognition of splice sites, and regulation of the competition between alternative sites, is aided by auxiliary regulatory elements that recruit splicing regulatory factors (SRFs).^5,6^ Finally, most splicing is thought to occur co-transcriptionally, with recognition of splice sites occurring relatively rapidly as RNA Polymerase II (RNAPII) is still transcribing.^7^ The accuracy of splicing decisions relies on the combined specificity of these genetic and biochemical interactions to assemble the spliceosome and catalyze intron excision, providing ample opportunities for stochastic interactions that might result in undesirable mRNA products.

Studies in the last few decades have started to investigate the error rate of splicing and quality control mechanisms to purge incorrectly spliced transcripts from the cell. Unlike DNA replication, and transcription or translation mechanisms, which have more narrowly defined error rates (10^-9^ basepairs and 10^-6^-10^-5^ nucleotides or amino acids, respectively^8–11^), less is known about splicing error rates (estimated to be between 10^-5^-10^-2^ junctions).^12–14^ Early quality control to minimize splicing error involves a process known as kinetic proofreading, which uses a family of DExD/H-box ATPases to outcompete suboptimal substrates and promote spliceosomal conformations for high fidelity splice site usage.^13,15,16^ It is thought that the spliceosome may iteratively sample potential splice sites and use kinetic proofreading to reject suboptimal sites before settling on the optimal site, similar to promoter proofreading by RNAPII.^17,18^ This process is not perfect, however, since there are many downstream “spell-checking” mechanisms to recognize and clear aberrantly or unspliced RNAs that escape earlier proofreading checkpoints.^16^ Spell-checking mechanisms degrade the undesirable substrate, either in the nucleus or cytoplasm. Nuclear turnover involves degradation by the exosome or XRN2, both of which compete with splicing on pre-mRNA.^15,19^ In the cytoplasm, turnover is often translation-dependent, with either nonsense mediated decay (NMD) or non-stop decay (NSD) triggered after the first round of ribosome translation.^20,21^

Despite extensive quality control steps to mitigate erroneous splice site usage, there is increasing evidence that “noisy” splicing regularly occurs at cryptic or low-fidelity sites. Previous studies have defined splicing noise in a number of ways, including (a) minor alternatively spliced isoforms from canonical splice sites,^2,22^ (b) unannotated cryptic splice sites that are used at very low frequency and/or are rapidly degraded,^12,23^ and (c) so-called low-fidelity splicing events that use non-consensus core dinucleotides.^14,23^ Additionally, non-canonical splice site usage can occur through recursive splicing, which is the use of juxtaposed 5′/3′ splice sites to trigger piecewise excision of long introns.^24,25^ Recursive splicing might be the result of stochastic interactions that actually aid in increasing splicing efficiency, especially in human cells,^26^ perhaps by reducing noisy splice site usage.^25^ The lack of a consistent definition for what constitutes noise in splicing has made it difficult to understand the full landscape of potential splicing interactions and pruning of noisy transcriptional output. Many studies have identified extensive non-canonical splicing of various forms across metazoan cells,^3,23,25,27–29^ but it is challenging to directly compare results across studies given the diversity of techniques applied to estimate non-canonical splice site usage. Thus, there is a lack of clarity on the intrinsic levels of noise and the ways in which cells are able to reduce the burden of this apparently wasted transcriptional output.^3^

We set out to answer a fundamental question in RNA biology: to what extent does stochastic splicing occur in mammalian cells? Here, using ultra-deep sequencing data to track splicing intermediates across RNA lifecycles, we systematically investigate the usage of cryptic and low-fidelity splice sites to understand noise in mRNA splicing. Our analyses reveal pervasive expression of cryptic sites in nascent RNA populations that does not persist to mature RNA, even after inhibition of translation-dependent degradation processes. We further show that while most cryptic splicing likely occurs stochastically, we can identify genomic features that can predict the occurrence or fate of a subset of these cryptic events.

## RESULTS

The selection of splice sites is thought to occur mostly co-transcriptionally, soon after sites become available on pre-mRNA transcripts.^30^ To maximize our ability to identify cryptic splicing events independent of their effects on transcript localization or stability, we selectively enriched for newly transcribed RNA. Specifically, we isolated nascent RNA from the nucleus after rapid metabolic labeling with 4-thio-uridine (4sU) for 8 minutes and pulling down 4sU-containing transcripts using a thiol-reactive biotin. We compared this nascent RNA with total nuclear RNA and, to track the ultimate fate of cryptic transcripts, polyA-selected RNA and polyA-selected RNA after cycloheximide (CHX) treatment to inhibit translation-mediated degradation mechanisms (e.g. NMD) (**Figure 1A**). We conducted these experiments independently in cells from K562 erythroleukemia and KNS60 glioblastoma cell lines and performed high-throughput sequencing on four replicates from each RNA enrichment method (except CHX-treatment, see Methods, **Table S1**). We validated the high quality of the RNA enrichment methods: (a) western blots of nuclear markers confirmed efficient nuclear fractionation **(Figure S1A)**; (b) nascent and nuclear samples showed higher levels of intronic reads derived from pre-mRNA molecules than seen in mature RNA **(Figure S1B)**, and (c) samples clustered by their RNA enrichment method across cell types (**Figures S1C**).

**Figure 1.**
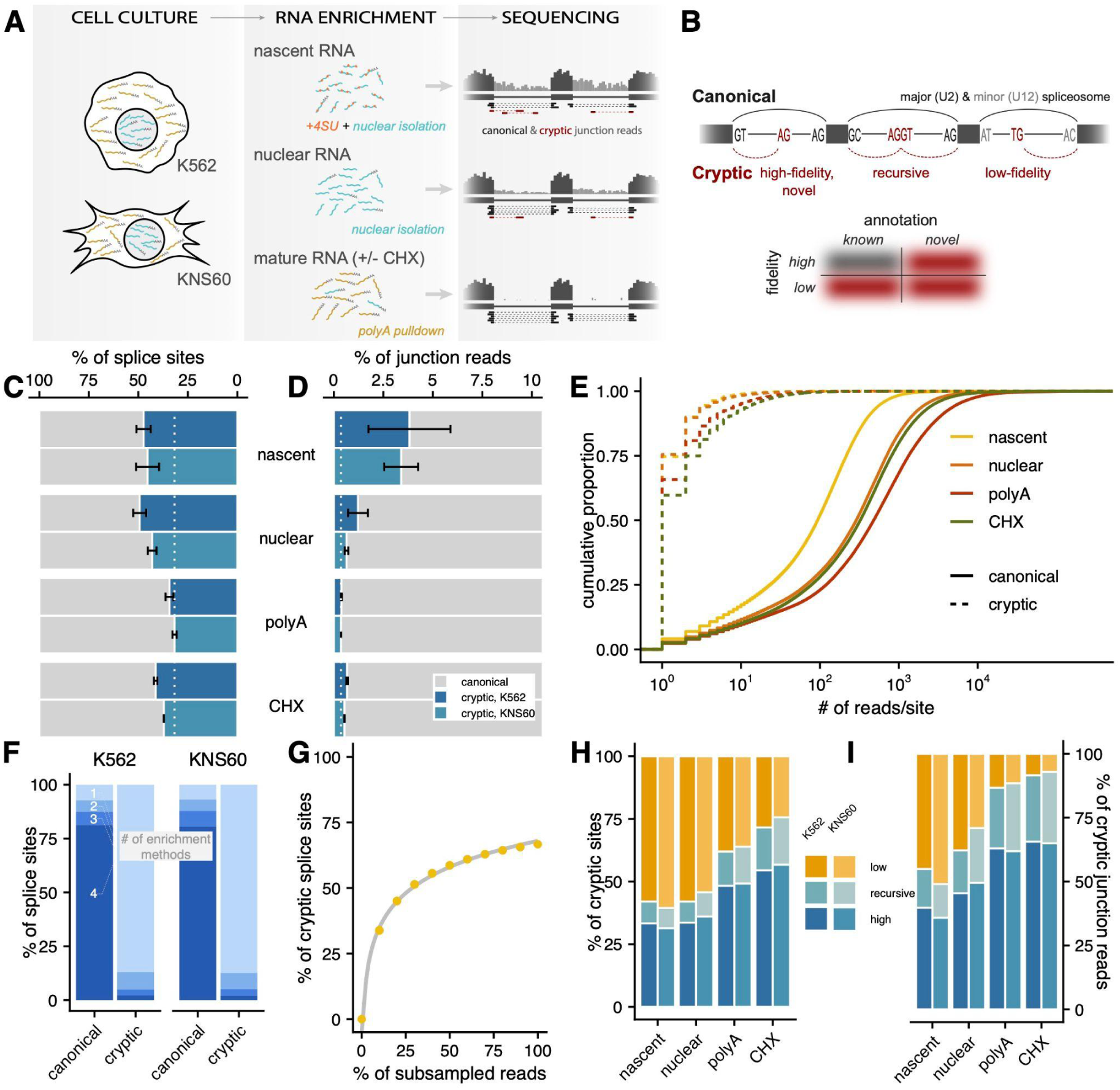
Cryptic splicing is pervasive across enrichment methods. **(A)** Schematic representation of the study design, RNA enrichment methods, and predicted patterns of high-throughput sequencing reads. **(B)** Schematic representation of types of canonical (*top*, *black*) and cryptic (*bottom*, *red*) splicing. Canonical dinucleotides typically recognized by the major and minor spliceosome are shown in black or grey, respectively, while cryptic dinucleotides are in red. **(C,D)** Percentage of canonical (*grey*) and cryptic (*blue*) splice sites and junction reads detected across enrichment methods and cell types. Dotted white line represents the mean percentage for polyA-enriched samples. **(E)** Cumulative proportion (*y-axis*) of junction reads per site (*x-axis*) for canonical (*solid*) and cryptic (*dashed*) splice sites across enrichment methods. **(F)** Percentage of canonical or cryptic splice sites that were detected in 1, 2, 3, or all 4 enrichment methods across cell types. **(G)** Percentage of cryptic sites (out of all splice sites, *y-axis*) detected across various sub-samples of read coverage (*x-axis*) in nascent RNA data, where 100% indicates the percentage of cryptic sites detected when reads from all four nascent RNA replicates are combined. **(H, I)** Percentage of cryptic sites and junction reads that can be classified as low-fidelity (*orange*), recursive (*teal*), or high-fidelity sites (*dark blue*) across cell types.

The position of splice sites used on pre-mRNA molecules can be inferred using high-throughput sequencing reads split across exon-exon boundaries (junction reads). However, technical or biological artefacts (e.g. overlapping reads, mapping irregularities, or sequence polymorphisms) may lead to mis-quantification of splicing events or split reads that are not derived from splicing events (**Figure S2A**). Thus, we developed a computational framework called CRYPTic IDentification of Splice Sites (*CRYPTID-SS*) to process high-throughput RNA-seq reads, identify high-confidence cryptic splicing events, and quantify their usage (Methods). To avoid known artefacts, we only considered uniquely mapped junction reads satisfying a number of criteria for further analyses, including those with sites within the same annotated gene boundaries and not overlapping polymorphisms or repeats (**Figure S2B**). Using these reads, we categorized canonical or cryptic splice sites based on the annotation status and sequence content of the core splice site dinucleotides. Canonical sites are defined as those present in the annotation databases and containing high-fidelity splice site motifs recognized by either the major or minor spliceosome, while cryptic sites are those that are either not included in annotation databases and/or containing low-fidelity dinucleotides (**Figure 1B**). Cryptic sites were further separated by whether the splice site dinucleotides indicated the probability of either a low-fidelity or high-fidelity binding event or supported the possibility of a recursive splicing event characterized by juxtaposed 3′ and 5′ splice site motif around a central AG|GT motif.

### Cryptic splicing is pervasive in nascent RNA

We identified 2,110,022 and 2,179,358 independent splice sites across all K562 and KNS60 samples, respectively, approximately evenly split between 5′ or 3′ splice sites, respectively. Using the cryptic vs. canonical definitions above, we observe a 41% increased number of cryptic splice sites, on average, in nascent and nuclear RNA than in mature RNA in both K562 and KNS60 cells (**Figure 1C**). This is complemented by a 206% increase, on average, in the number of junction reads that support the use of cryptic splice sites in nascent and nuclear samples compared to mature RNA (**Figure 1D**). However, while canonical sites are supported by more than 100 reads on average across all enrichment conditions, cryptic sites rarely have more than 1 supporting read (**Figure 1E**), indicating that splicing at any given cryptic site occurs very infrequently. Concordantly, 87% of cryptic sites are detected in only one enrichment method (**Figure 1F**) — in contrast to canonical sites, most of which are detected in all four enrichment methods — and an average of 84% are present in only one replicate from that enrichment method (**Figure S3A**). We assessed the sensitivity of our detection pipeline by running it on subsamples of reads ranging from 10% to 100% of the total reads collected for each enrichment method. New cryptic sites were still being detected as read depth increased from 50% to 100% of sequenced reads (**Figure 1G**) and thus would likely increase further at higher read depths. However, the rate at which new sites are detected starts to plateau in this range, suggesting that there is an upper limit on the number of genomic positions that can be used as cryptic splice sites.

To rule out the possibility that the pervasive detection of rarely used cryptic splice sites was not due to technical artefacts, we performed the same analyses with a series of control datasets. First, to examine how splice site usage is affected by 4sU treatment, we used polyA RNA from cells treated with uridine and 4sU for 8 minutes or 24 hours and see no substantial increase in the number of cryptic sites that are expressed (**Figure S3B,C**). Second, to examine the influence of mapping errors on the identification of cryptic sites, we identified “cryptic splice sites” in an DNA exome sequencing dataset from K562 cells where all junction reads should be the result of sequence variation or mapping biases. We see that on average, 0.9% of splice sites in K562 RNAseq were also observed in the K562 exome data and we removed these sites from the final analyses. Finally, previous reports have suggested that library preparation may induce spurious junction reads (*e.g.* through trans-splicing^31^). Using both oligo-dT and random hexamer primed libraries from synthetic Spike-In RNA Variant (SIRV) mixes, we see that less than 0.3% of junction reads are erroneous (**Figure S3D,E**). Since SIRVs provide a ground-truth for the fraction of erroneous junctions expected in the sequencing data, we also used these data to define a false discovery rate (Methods) and estimate that 0.06% of our cryptic sites, after the filtering steps above, are expected to be false positives.

Previous methods to identify cryptic splice sites have hypothesized that most cryptic molecules are degraded by mechanisms such as NMD so blocking translation should cause those molecules to be retained in a mature RNA population.^27,29,32^ While we do see a higher percentage of cryptic sites and reads in RNA from CHX treated cells (19% and 61% increases, respectively, **Figure 1C,D**), these increases are modest relative to those seen in nascent RNA. Treatment with a chemical NMD inhibitor (NMDi14) gave consistent results with CHX treatment, while a DMSO control showed no increase in cryptic sites (**Figure S3B,C**). When we sub-categorize cryptic sites, we see that the majority of cryptic sites in nascent RNA are low-fidelity while the majority of cryptic sites in the CHX-treated RNA are high-fidelity (**Figure 1H**), though low-fidelity sites are generally supported by the fewest reads (**Figure S3F**). Overall, the extent of low-fidelity splicing gradually decreases as RNA becomes more mature (**Figure 1H,I**), suggesting that molecules with these splicing events are even less stable than products of other types of cryptic splicing. Intriguingly, the proportion of sites classified as recursive sites increase as RNA matures, which is contrary to the expectation that these sites would be spliced out when their parent intron is fully spliced out and suggests that splicing at recursive motifs could also result in stable mRNA products. Finally, the increase in the extent of high-fidelity cryptic splicing in RNA from CHX-treated cells suggests that these sites are the most likely to result in NMD. Overall, these results indicate that there is pervasive cryptic splicing — including at motifs that the spliceosome is not thought to bind — and that the perturbation of translation-dependent degradation mechanisms does not completely uncover the full landscape of splice site usage on pre-mRNA molecules.

### Cryptic splicing occurs throughout genes

Our expanded catalog of cryptic splice sites across RNA life stages (nascent RNA, nuclear RNA, mature RNA, and NMD-sensitive RNAs) enables a systematic assessment of the regulatory environments in which cryptic splicing occurs. To do so, we focused on splice sites that had support from at least two uniquely mapping reads, were detected in at least two independent replicates, or both (**Table S2**). Using this set of high-confidence sites, we first looked at the distribution of sites across a gene. Canonical 5′ splice sites are distributed in a hockey stick-like distribution (**Figure 2A**) across a gene, with a peak of sites at the 5′ end of a gene followed by a dip and slow increase of sites that is likely driven by first introns often being the longest intron in human genes.^33^ Correspondingly, canonical 3′ splice sites steadily increase across the gene body. In contrast, cryptic sites in nascent and nuclear RNA are much more evenly distributed across the gene, with both 5′ and 3′ cryptic sites slightly more likely to occur near the beginning of the gene (**Figure 2A; Figure S4A**), potentially reflecting their placement in the first intron. We see that the distribution of cryptic sites in nascent and nuclear RNA closely matches the distribution of nucleotides across UTRs, coding exons, or intronic regions (**Figure 2B; Figure S4B**), with a modest enrichment of recursive and high-fidelity sites in coding exons (Odds Ratio = 1.56-2.56; Hypergeometric Test p-value < 2.2 x 10^-16^ for both cryptic types). In mature RNA, the distribution of cryptic sites more closely matches the distribution of canonical sites and they are significantly are depleted in introns (Odds Ratio = 0.14; Hypergeometric Test p-value < 2.2 x 10^-16^), with low-fidelity sites sharply enriched at the 3′ end of the gene and in 3′ UTR regions (Odd Ratio = 14.97; Hypergeometric Test p-value < 2.2 x 10^-16^; **Figure 2A,B; Figure S4A,B**). These observations suggest that while there is promiscuous spliceosomal binding and function across a gene, the positioning of cryptic sites (particularly in 3′ UTRs) likely influences transcript stability or the potential to trigger NMD.

**Figure 2.**
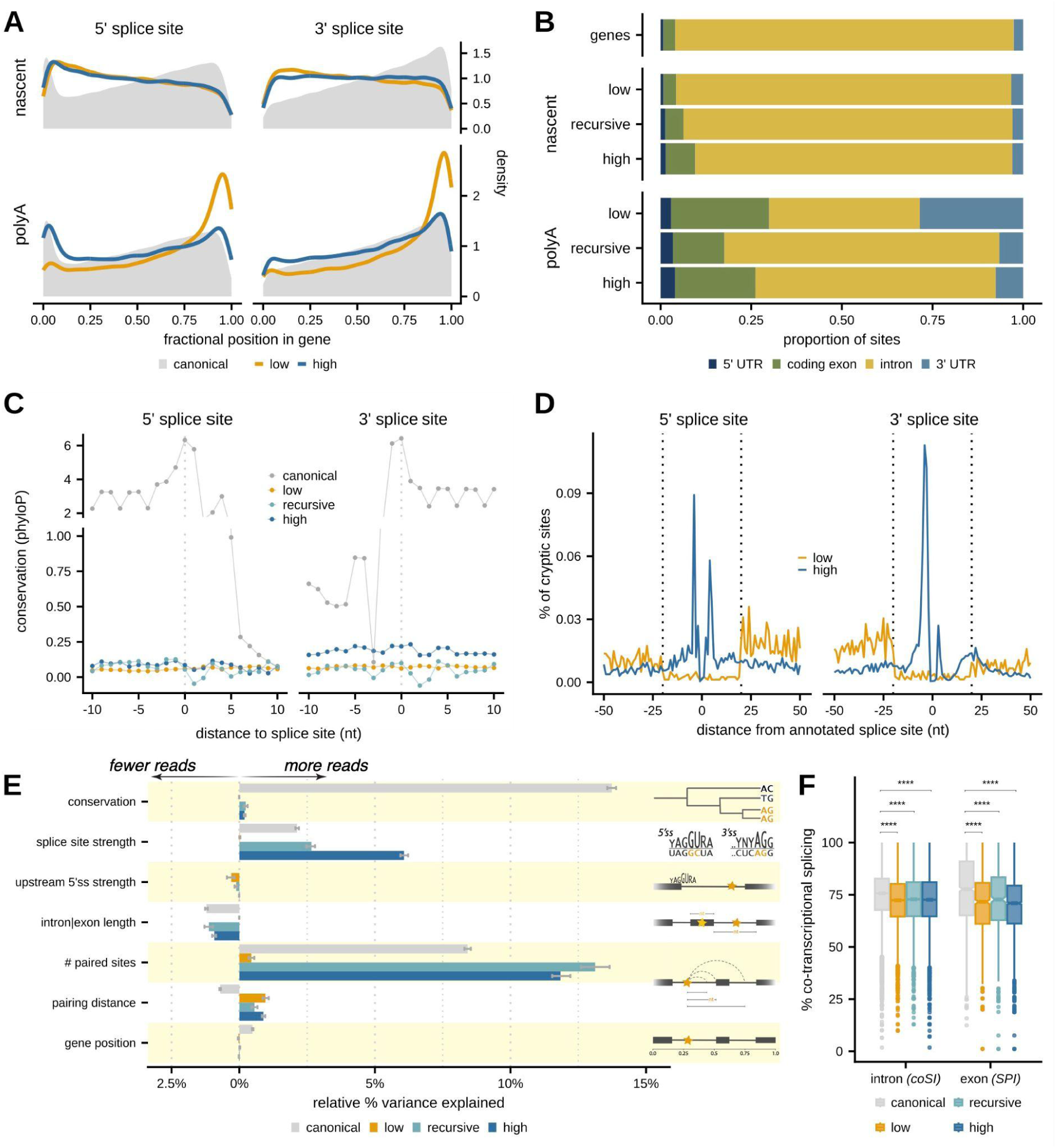
Genomic features influence the use of cryptic sites. **(A)** Distribution of the position of 5′ (*left*) and 3′ (*right*) splice sites across a gene categorized as canonical (*grey*), low-fidelity (*orange*), or high-fidelity (*dark blue)* in either nascent (*top*) or mature (*bottom*) RNA, where the fractional position is defined as the distance of a site from the start of a gene relative to the total gene length. **(B)** The proportion of each type of cryptic site detected within annotated 5′ UTRs (*navy blue*), coding exons (*green*), introns (*yellow*), and 3′ UTRs (*light blue*) for sites detected in nascent (*middle*) or mature (*bottom*) RNA. The top bar shows the proportion of nucleotides within expressed genes for each annotation category. **(C)** Nucleotide-specific phyloP scores (*y-axis*) in a 20nt window around 5′ and 3′ splice sites categorized as canonical (*grey*), low-fidelity (*orange*), recursive (*teal*), or high-fidelity (*dark blue*) sites. **(D)** Distribution of the distance that a cryptic 5′ or 3′ splice site occurs relative to the nearest canonical 5′ or 3′ splice site, respectively, for low-fidelity (*orange*) or high-fidelity (*dark blue*) sites. **(E)** Relative importance of features influencing variance in splice site usage for canonical (*grey*), low-fidelity (*orange*), recursive (*teal*), and high-fidelity (*dark blue*) sites, using multiple linear regression models. Bars pointing to the right indicate positively correlated features, while bars pointing left indicate negatively correlated features. Schematics on the right provide a visual representation of each feature. **(F)** The distribution of completed splicing index (coSI, *left*) or splicing per intron (SPI, *right*) for splice sites in annotated introns or exons, respectively, categorized by canonical (*grey*), low-fidelity (*orange*), recursive (*teal*), or high-fidelity (*dark blue*) sites. Signficance was assessed with a Mann-Whitney U Test. **** adjusted p-value < 0.0001.

Splice sites are considered to be among the most conserved sites in mammalian genomes, given their crucial regulatory roles for establishing the sequences of mRNA and proteins. However, if cryptic sites reflect noisy, rather than regulated, splicing, they might not show the same conservation patterns as canonical sites. We see that the core dinucleotides for canonical 5′ and 3′ splice sites show extremely high conservation (**Figure 2C**), with sustained conservation into exonic regions that exhibits triplet periodicity reflective of less conservation of the 3rd “wobble” nucleotide in a codon.^23^ However, cryptic sites exhibit little to no signature of conservation, though there is weak evidence for triplet periodicity downstream of 3′ cryptic recursive or high-fidelity sites. Interestingly, while other studies have seen moderate to strong higher conservation for bona fide recursive splice sites in mammalian and particularly in fly cells,^24,25,34–36^ the recursive sites that we classify based on motifs alone show no increase in conservation relative to low-fidelity cryptic sites. The pattern of slightly higher conservation around high-fidelity 3′ splice sites led us to speculate that this might reflect NAGNAG alternative splicing, which is an alternative 3′ splice site mechanism that produces two or more alternative isoforms differentiated by three nucleotides (NAG) at the 5′ end of an exon.^37^ Thus, we looked at the distribution of cryptic sites around annotated splice sites. While ∼94% of overall cryptic splicing occurs more than 50 nucleotides from an annotated splice site, we see that high-fidelity cryptic sites are enriched in the 25 nucleotides upstream and downstream of annotated sites (**Figure 2D**). This is particularly true for 3′ splice sites, where we observe that as many as 10% of high-fidelity 3′ sites might be NAGNAGs alternative splice sites. In contrast, both 5′ and 3′ low-fidelity sites appear to be almost completely depleted in the same 25nt footprint around annotated sites and their density increases approximately 25nt into intronic regions. While this might be related to a footprint of strong spliceosome binding precluding cryptic activity near canonical sites, it is also possible that the slight enrichment of high-fidelity cryptic sites inherently leads to a depletion of low-fidelity dinucleotides in this region.

Together, our results thus far suggest that cryptic sites (especially low-fidelity sites) are used at random across a gene. To directly test this, we used a multiple linear regression framework to estimate the contributions of genomic features to the usage of individual cryptic sites (Methods). We fit this model separately to each type of cryptic site across enrichment methods and find that this framework can explain 22% of variance in canonical site usage and only 2-17% of variance in cryptic site usage in nascent RNA (**Figure 2E**). Unsurprisingly, the usage of canonical sites was positively associated with the evolutionary conservation and strength of the sites. Splice site strength also explained 3-5% of variance in the usage of recursive and high-fidelity sites (particularly in mature RNA; **Figure S4C**), which is consistent with the usage of these sites being driven by the increased probability of spliceosomal binding due to similarity with canonical splice sites. Interestingly, the number of other sites that a cryptic site paired with was one of the strongest contributors to splice site usage, such that sites were more likely to be used if they were paired with more partner sites that were farther away. This relationship was true for cryptic sites even when correcting for the number of partner sites (**Figure S4D**). Furthermore, low-fidelity splice sites in particular were also less likely to be used when located downstream of a strong canonical 5′ splice site, suggesting that stronger 5′ splice sites might lead to faster splice site binding and commitment to a splicing reaction that otherwise might involve a cryptic site. Consistent with this, cryptic splice sites were more likely to fall into introns or exons that show less evidence for co-transcriptional splicing (**Figure 2F**). These observations point to the local splicing environment being an important contributor to cryptic splice site usage, such that inefficient splicing of canonical sites might lead to an opening for cryptic sites to pair with a partner site and complete splicing after promiscuous binding of the spliceosome.

### Sequence determinants of low-fidelity splicing

While 5′ and 3′ splice sites are likely recognized as independent units, the favorable pairing of two sites across the intron is necessary for splicing to occur (**Figure 3A**). Our observation that site pairing contributed to cryptic site usage led us to further investigate pairing properties between 5′ and 3′ splice sites. Across RNA life stages, we see that the majority of pairs involve two canonical sites (**Figure 3B**), while 57% of cryptic sites are paired with other cryptic sites in nascent RNA. On average, canonical sites pair with a site that is slightly more distant than the closest annotated site, while cryptic sites often pair with sites that are closer than the closest annotated site (**Figure S5A**). The proportion of cryptic-cryptic site pairing decreases as RNA matures, with 83-87% of cryptic sites in mature RNA paired with canonical sites, indicating that transcripts with cryptic splicing are more stable or have a higher probability of surviving to maturity when cryptic sites paired with canonical sites. The increase in pairing of cryptic sites with canonical sites across RNA life cycles mostly reflects the pairing propensities of recursive and high-fidelity sites, while almost all low-fidelity sites are paired with other cryptic sites (**Figure 3C, Figure S5B**). Interestingly, there is more pairing between recursive 3′ and 5′ sites in both nascent and mature RNA (**Figure S5B)**, while recursive 5′ sites are more likely to pair with canonical sites. This indicates that recursive splicing most likely does not occur in a sequential (upstream-segment-first) order across an intron, but rather, consistent with recent reports^26^, individual segments flanked by recursive sites can be spliced out of larger introns.

**Figure 3.**
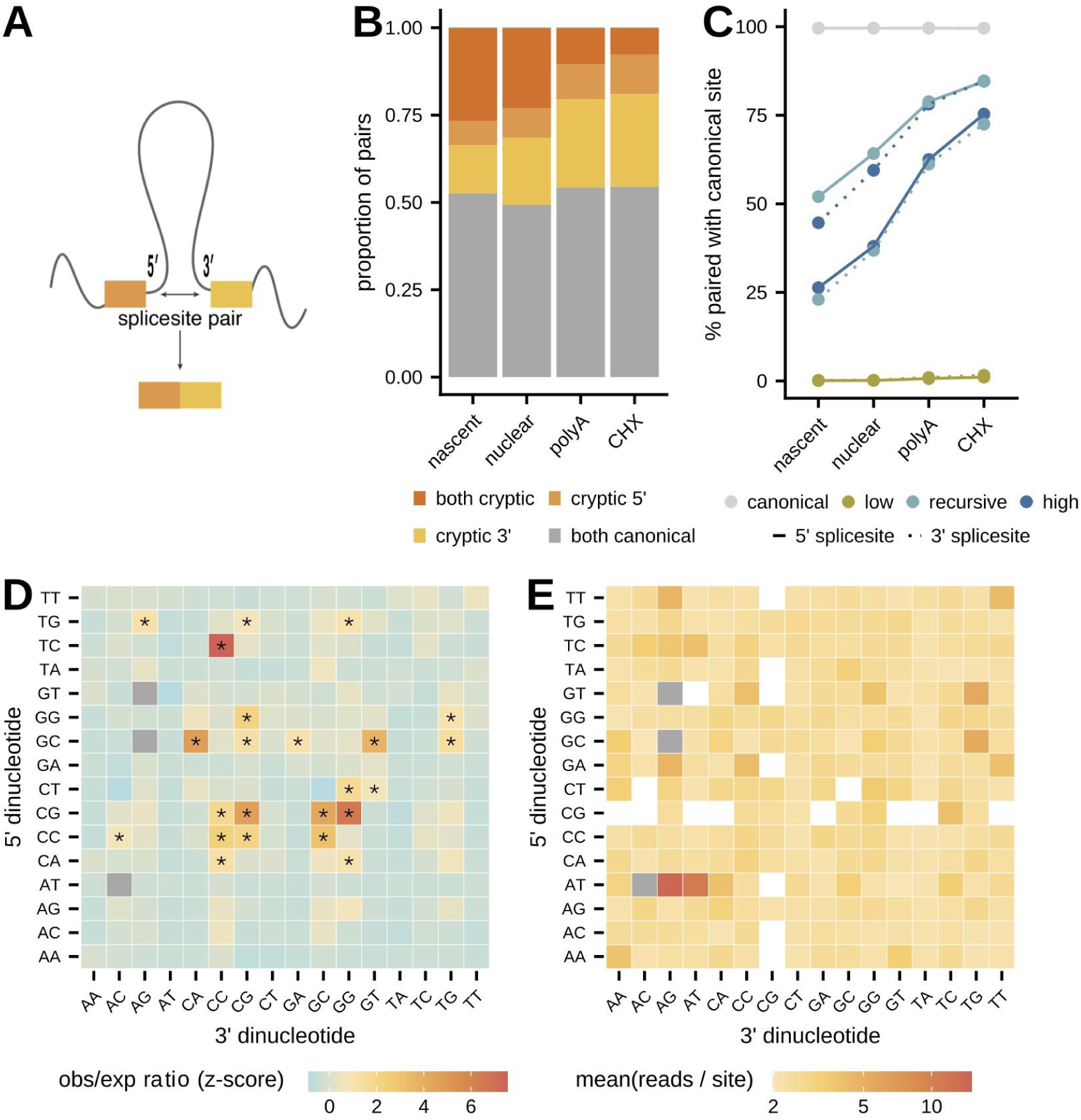
The pairing of low-fidelity splice sites is driven by specific dinucleotide preferences. **(A)** Schematic representation of pairing between 5′ and 3′ splice sites that define an intron to be excised. **(B)** Proportion of splice site pairs in which both, only one, or neither of the splice sites are cryptic, across enrichment methods. **(C)** Percentage of 5′ (*solid line*) or 3′ (*dashed line*) canonical (*grey*), low-fidelity (*orange*), recursive (*teal*), or high-fidelity (*dark blue*) sites that are paired with canonical sites across enrichment methods. **(D)** Heatmap of the ratio between the observed number of sites relative to the expected distribution of sites (calculated by estimating the frequency each dinucleotide occurs in expressed genes) for each combination of 5′ (*y-axis*) and 3′ (*x-axis*) dinucleotides used by low-fidelity splice sites detected in nascent RNA. Dinucleotide pairs involved in canonical splice site pairings were omitted from the plot (*grey boxes*). Significance was assessed using permutations. * adjusted p-value < 0.05. **(E)** Heatmap of the mean number of reads per site for each combination of 5′ (*y-axis*) and 3′ (*x-axis*) dinucleotides used by low-fidelity splice sites detected in nascent RNA. Dinucleotide pairs involved in canonical splice site pairings and pairs with fewer than 25 observations were omitted from the plot (*grey* and *white boxes*, respectively).

These observations suggested that there might be sequence preferences that underlie the probability of successful pairing and splicing involving cryptic sites. Overall, we do not see any particular sequence contexts within which cryptic sites are more likely to occur and do not see enrichments of any particular dinucleotides among low-fidelity sites. Instead, we see that there are 23 pairs of dinucleotides that are significantly enriched with 2.19 fold more sites, on average, than expected relative to their joint occurrence in transcribed regions (**Figure 3D**, Methods). Interestingly, 91% of the preferred dinucleotide pairs are GC-rich, suggesting that local sequence content, mRNA secondary structure, or stability might play a role in the usage of these sites. Of note, it has been seen that G-quadruplex (G4) motifs are enriched near splicing junctions and the formation G4 structures enhance splice site usage.^38^ However, these overrepresented sequence pairs do not represent the most frequently expressed cryptic junctions. Instead, the dinucleotide pairs that are used the most are AT-AG and AT-AT (**Figure 3E**), suggesting that the minor spliceosome may be more likely to use different 3′ splice sites than previously thought.^39^ Importantly, we do not see similar enrichments in the control dataset generated from synthetic SIRVs (**Figure S5C,D**), indicating that these enrichments are not likely due to errors or biases that occur during library preparation, sequencing, or mapping.

### Longer, highly expressed genes have more cryptic splicing

Our results suggest there is pervasive cryptic splicing driven primarily by stochastic spliceosome binding across pre-mRNA transcripts. If this were true, cryptic splicing would be more likely to occur in genes whose genomic or transcriptional properties presented more opportunity for splicing noise. For instance, longer genes present greater pre-mRNA real estate for spliceosomal binding at random. Similarly, if every pre-mRNA transcript has the same probability of having a cryptic splicing event, high transcription levels would result in a greater occurance of cryptic splicing in the gene. However, mRNA transcript composition is known to be highly regulated and under strong evolutionary constraints to ensure proper mRNA or protein regulation. Thus, we wondered how variable cryptic splicing was across genes with different genomic features and if genes varied in their tolerance for cryptic splicing.

To do so, we first quantified the proportion of transcripts from each gene that had cryptic splice sites. We devised a junction TPM, which calculates the relative expression of each type of cryptic transcript using only empirical counts of exon-exon junction reads (Methods). We ranked genes by these junction-derived TPMs and see that there is large variability in the extent of cryptic splicing per gene. Overall, the proportion of cryptic transcripts is negatively correlated with both transcription levels and steady-state gene expression levels (assessed from nascent RNA and mature RNA, respectively; Methods) and number of introns in a gene, but positively correlated with the length of the gene and fraction of a gene annotated to be intronic (**Figure 4A; Table S3**). Genes appear to be clustered into 3 distinct categories: (1) 540 (4.2%) genes with no cryptic splicing, which tend to be lowly transcribed, moderately expressed, small genes that are enriched for genes involved in oxygen binding and gas transport, (2) 11,071 genes (87%) with cryptic transcripts accounting for anywhere from 1-99% of their total transcripts, and (3) 1,053 (8.3%) genes with nearly or exactly 100% cryptic transcripts — including 424 genes with 100% cryptic splicing but no low-fidelity cryptic splicing, which tend to be lowly transcribed, lowly expressed genes with few annotated introns. Overall, this last category is enriched for genes involved in olfaction, structural components of chromatin (*i.e.* histone genes), and electron transport chain activity, suggesting that regulatory pressures on these pathways might be reduced when these genes are expressed at low levels.

**Figure 4.**
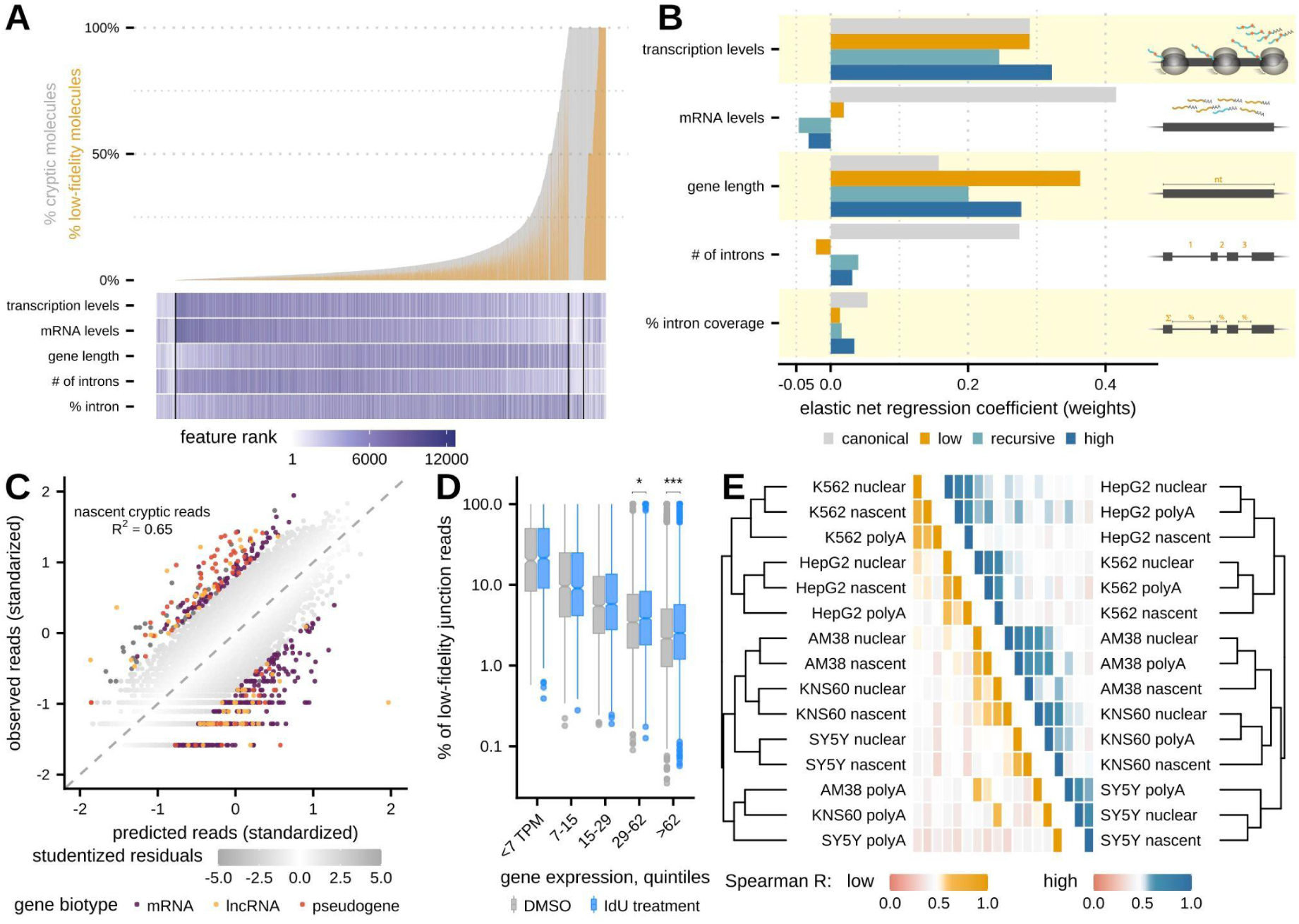
Highly expressed and longer genes are more prone to cryptic splicing. **(A)** The ranked distribution of the percentage of transcripts with any cryptic splicing (*grey*) and specifically with low-fidelity splicing (*orange*) per gene, using junction-specific TPM estimates. Below are the associated transcription levels (nascent RNA TPMs), mRNA levels (mature RNA TPMs), gene length (nucleotides), number of total introns, and the percent of a gene sequence that is annotated as intronic, where each color indicates the rank of a feature ordered from lowest value to highest value. **(B)** Feature weights from elastic net regression models (*x-axis*) trained on the number of cryptic splicing reads per gene for canonical (*grey*), low-fidelity (*orange*), recursive (*teal*), and high-fidelity (*dark blue*) sites in nascent RNA. Schematics on the right provide visual representations of each feature, which were estimated as described in A. **(C)** The amount of cryptic splicing per gene predicted by the elastic net regression model (center standardized, *x-axis*) versus the observed cryptic splicing per gene in nascent RNA (*y-axis*). The grey color scale indicates the magnitude of studentized residuals and genes with |studentized residual| ≥ 2 are colored by their ENSEMBL biotype. **(D)** Distribution of the percentage of low-fidelity junction reads (relative to all junction reads) per gene (*y-axis*) in control (*grey*) and IdU-treated cells (*blue*) for genes in different bins of gene expression levels (*x-axis*). Signficance was assessed with a Mann-Whitney U Test. * adjusted p-value < 0.05. *** adjusted p-value < 0.001 **(E)** Clustering of samples across cell types and enrichment methods by the extent of low-fidelity (*left*) or high-fidelity (*right*) splice sites per gene. Dendrograms indicate hierarchical clustering distances, while heatmaps indicate pairwise Spearman correlation values.

While there are clear associations between the extent of cryptic splicing and genomic features of a gene, the high degree of correlations between pairs of these features makes it difficult to disambiguate the relative role of each in influencing the likelihood of cryptic splicing for a gene. To overcome this challenge, we trained an elastic net regression model, which specifically accounts for correlated variables, to predict the gene-specific usage of cryptic splice sites based on the combined influence of these genomic features (Methods). Overall, we can explain 58-81% variance in both the number of cryptic sites and the number of cryptic reads for a gene across all types of splice sites for nascent and nuclear RNA (**Figure S6A**). The features that contributed the most to the number of cryptic events and the usage of sites in nascent RNA were transcription levels and gene length, both positively (**Figure 4B**; **Figure S6B**). Low-fidelity sites are particularly associated with longer gene lengths. Higher mRNA levels were weakly negatively associated only with recursive and high-fidelity splicing, as expected if those transcripts were more likely to be degraded after reaching maturity. In contrast, these features only explain 44-74% of variance across cryptic sites in mature RNA, while mRNA levels, with the number of introns explaining 64% of variance in the canonical splice site usage (**Figure S6C**).

We used our predictive framework to identify genes with more or less cryptic splicing in nascent RNA than expected given their genomic features. Using a conservative threshold to define outliers, we see that more genes (3.8%) have less cryptic splicing than expected and that 81% of these genes are mRNAs (**Figure 4C**). However, only 1.6% of genes have more cryptic splicing than expected and these genes are enriched for expressed pseudogenes or lncRNAs (39% of genes; Odds Ratio = 6.93; Hypergeometric Test p-value = < 2.2 x 10^-16^). This is consistent with the idea that these types of RNAs might have less splicing regulation and/or less selective constraints on their sequence content,^40,41^ leading to an increased tolerance for cryptic splicing without severe consequences for any functional role of the lncRNA. It has been hypothesized that lncRNA-spliceosome complexes themselves — rather than fully spliced lncRNA — might serve a role in stimulating local gene expression, suggesting that functional splicing on lncRNAs might be agnostic to the specific position at which it occurs.^40^ In contrast, approximately equal numbers of genes (∼2-3%) have more or less cryptic splicing than expected in mature and CHX-treated RNA (**Figure S6D**). Genes with less cryptic splicing in mature RNA are enriched for genes involved in cytoplasmic translation, including those that encode for ribosomal protein genes.

### Transcriptional variation influences cryptic splicing

As expected if cryptic splicing was mostly stochastic, the features most strongly associated with cryptic splicing in a gene increased opportunities for splicing to occur. In light of this, we wondered if cryptic splicing was more generally driven by mechanisms that led to more transcriptional noise. To test this, we treated K562 cells with 5′-iodo-2′-deoxyuridine (IdU) — a small molecule that has been shown to increase transcriptional variance without affecting the mean of gene expression^42^ — and collected nascent RNA. For genes that are more highly expressed and thus allow for more cryptic site detection, we see significant increases in cryptic splice site usage, particularly among low-fidelity splice sites (**Figure 4D**). If transcriptional fluctuations were a primary driver of cryptic splicing across genes, we would also expect that genes should have similar extents of cryptic splicing across cell types after normalizing for RNA abundance. To evaluate this, we generated nascent, nuclear, and mature RNA data from three additional cell lines — hepatocellular carcinoma cells (HepG2), neuroblastoma cells (SY5Y), and an additional glioblastoma line (AM38) — and evaluated the extent of cryptic splicing across cells after normalizing for RNA abundance within the RNA enrichment method (Methods). Samples tend to cluster first by their cell type of origin and then RNA life stage (**Figure 4E**), both when clustering by low- and high-fidelity splicing. The two glioblastoma cell types (KNS60 and AM38) cluster together, following clustering of all the neuronally derived cell types. Of note, the mature RNA samples from the neuronally derived cell types cluster apart from the nascent and nuclear samples, perhaps due to the relative lack of low-fidelity splicing that persists in mature RNA. These results indicate that while there is abundant evidence for stochastic usage of cryptic sites, there are differences in either the propensity for cryptic splicing or gene-specific tolerance for cryptic splicing across cell types.

### Molecular determinants of cryptic splicing

In order to further understand molecular influences on cryptic splicing levels, we sought to identify RNA-binding proteins (RBPs) that regulate global cryptic splicing levels in cells. To do so, we used RNA-seq data from siRNA-mediated knockdown of 204 RBPs from the ENCODE consortium^43^ and used *CRYPTID-SS* to evaluate significant changes in junction reads observed in the knockdown (KD) relative to a batch-matched scrambled siRNA control (Methods). We see that the majority of RBP KDs result in global shifts towards more recursive (68% of factors) and/or high-fidelity (75%) reads (**Figure 5A**) or sites (**Figure S7A**), with 4 factors resulting in a significant increase in recursive and high-fidelity cryptic splicing: SUPV3L1, EIF4A3, HNRNPC, and AQR. Each of these factors is known to play a role in splicing regulation or quality control: the mitochondrial (mt) RNA helicase SUPV3L1 is involved in mtRNA processing and degradation,^44^ DEAD-box RNA helicase EIF4A3 is a core component of the exon junction complex (EJC) that plays a role in splicing and NMD,^45^ and HNRNPC functions as a molecular ruler to measure RNA transcript length before nuclear export^46^ and is a reader of m6A RNA modifications.^47,48^ Finally, AQR is an RNA helicase that is a component of the intron binding complex^49^ and is required for efficient pre-mRNA splicing by potentially disassembling spliceosomes on poor substrates.^50–52^ Our results are consistent with previous studies that have identified AQR, and to a lesser extent EIF4A3, as regulators of cryptic high-fidelity splice site usage.^53–55^ In contrast, the majority of factors result in shifts towards slightly less canonical and/or low-fidelity splicing, with no factors that significantly change the levels of low-fidelity splicing. However, since these data are derived only from mature RNA, we are unable to distinguish between factors that may change the propensity for cryptic splice sites to be used and factors that are involved in the quality control or stability of transcripts with cryptic sites.

**Figure 5.**
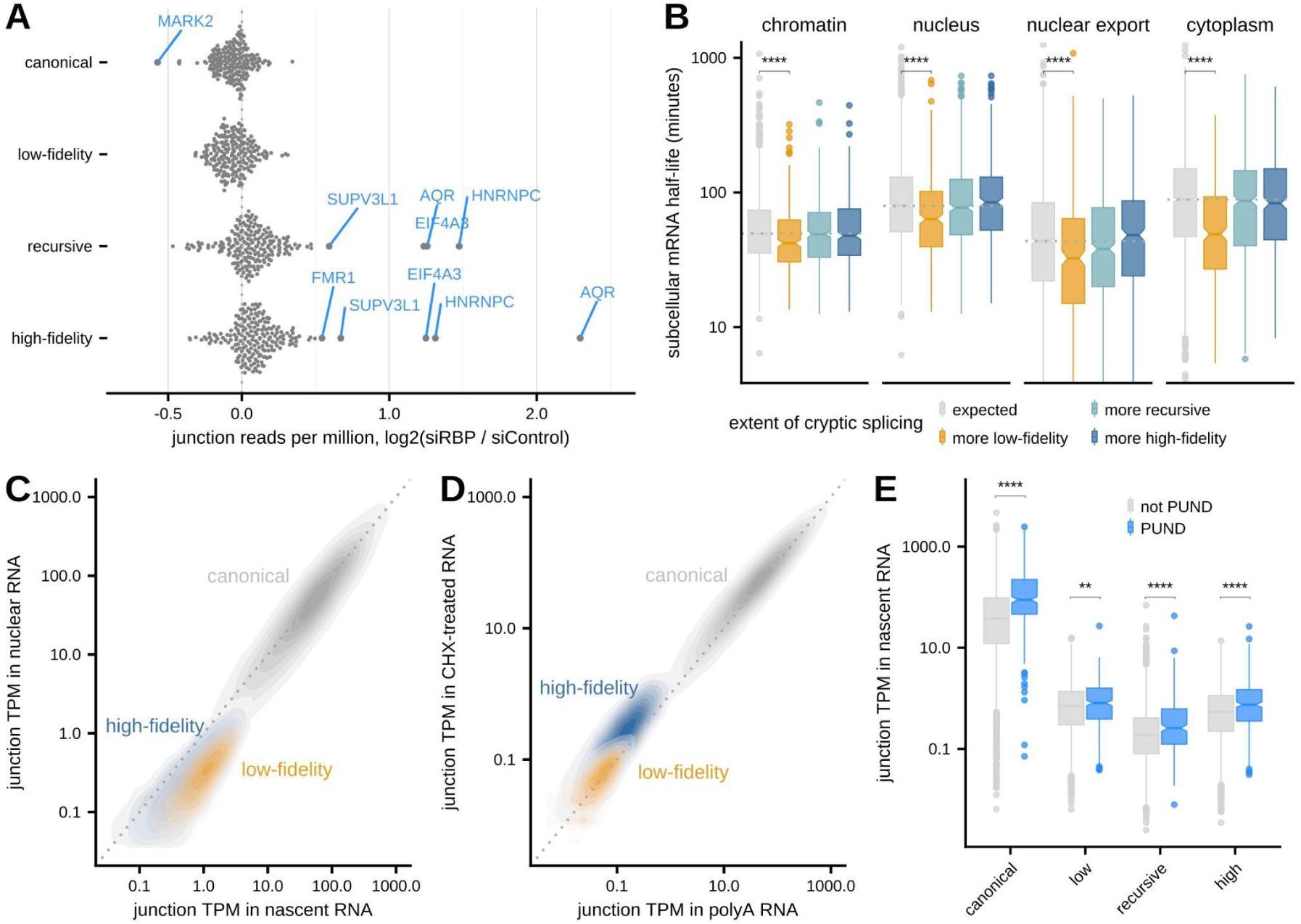
Cryptic transcripts are quickly cleared out of cellular compartments. **(A)** The fold change in junction reads between RNA binding protein siRNA knockdown and scrambled siRNA control samples (*x-axis*) for canonical, low-fidelity, recursive, and high-fidelity junction reads across all RBPs assessed by the ENCODE consortium. Significance was estimated using a permuted null distribution using control samples and significantly changing factors are labeled in blue (adjusted p-value < 0.05). **(B)** Distributions of mRNA residence times or half-lives across different cellular compartments estimated using subcellular TimeLapse-seq^56^ for genes with more low-fidelity (*orange*), recursive (*teal*), or high-fidelity (*dark blue*) transcripts than predicted by the elastic net regression model, with genes whose cryptic splicing levels match expected levels shown in grey. Signficance was assessed with a Mann-Whitney U Test. **** adjusted p-value < 0.0001. **(C)** Scatterplot of junction-specific TPMs in nascent RNA (*x-axis*) versus nuclear RNA (*y-axis*), separated by canonical (*grey*), low-fidelity (*orange*), and high-fidelity (*dark blue*) transcripts per gene. **(D)** Scatterplot of junction-specific TPMs in mature RNA (*x-axis*) versus CHX-treated mature RNA (*y-axis*), separated by canonical (*grey*), low-fidelity (*orange*), and high-fidelity (*dark blue*) transcripts per gene. **(E)** The distribution of junction-specific TPMs (y-axis) across splice site types, separated by genes categorized as predicted to undergo nuclear degradation (PUNDs^56^, *blue*) or matched controls (*grey*). Signficance was assessed with a Mann-Whitney U Test. ** adjusted p-vale < 0.01. **** adjusted p-value < 0.0001.

### The fate of cryptic transcripts depends on the splice site fidelity

Finally, we wanted to understand the ultimate cellular fate of transcripts with cryptic sites. Our initial observations suggested that transcripts with cryptic sites — in particular, those with low-fidelity sites — are likely pruned out from nuclear and mature RNA pools by quality control or degradation mechanisms. While high-fidelity cryptic splicing shows similar properties through RNA enrichments and is more abundant after CHX treatment, low-fidelity sites substantially decrease in frequency through RNA life stages. To evaluate the lifetime of cryptic transcripts and the cellular compartments in which they might be subject to regulation, we used subcellular residence times in K562 cells estimated using the recent RNAFlow approach from Ietswaart *et al.*^56^ We see that genes with more low-fidelity splicing than expected (relative to elastic net regression model predictions) spend less time in each compartment and have shorter half-lives. In particular, they spend significantly less time on chromatin, less time in the nucleus, have faster nuclear export rates, and are degraded faster in the cytoplasm (**Figure 5B**). One possibility is that these transcripts could be generally unstable in every cellular compartment. However, it is also possible that transcripts using low-fidelity splice sites are recognized co-transcriptionally (leading to faster chromatin release, **Figure S7B**), quality control mechanisms lead to nuclear degradation, and thus there are fewer of these transcripts to export and degrade in the cytoplasm.

While it is challenging to distinguish between these two possibilities directly, we used the relative expression of cryptic transcripts across cellular compartments to indirectly ask about where degradation might be occurring. We find that transcripts with low-fidelity splice sites are expressed 3.3 fold more on average in nascent RNA than in the overall nuclear RNA population, while the expression of high-fidelity cryptically spliced transcripts is less shifted (2.4 fold) and transcripts with canonical sites are highly correlated between the two factions (Pearson R = 0.91; 1.1 fold more in nascent RNA; **Figure 5C**). However, when looking at mature RNA, transcripts with high-fidelity splicing are much more affected by transcription inhibition with CHX (1.77 fold more expressed in CHX-treated RNA on average) than the expression of transcripts with canonical or low-fidelity sites (Pearson R = 0.97 and 0.96, respectively; both 1.1 fold more in CHX-treated RNA; **Figure 5D**). Together, these results suggest that many transcripts with cryptic sites, and especially those with low-fidelity splicing, are recognized and degraded in the nucleus, while transcripts with high-fidelity splicing are more likely to survive long enough to trigger cytoplasmic translation-dependent degradation mechanisms. Consistent with this, we see that the 340 genes predicted to undergo nuclear degradation (PUNDs) by RNAflow analyses^56^ have significantly more transcripts with cryptic splicing in nascent RNA than non-PUND genes (**Figure 5E**). However, PUNDs also have more expression of transcripts with canonical splicing as well, perhaps pointing to greater splicing activity overall in this class of genes.

## DISCUSSION

Here, we systematically identify and quantify cryptic splicing across stages of the RNA lifecycle. Our results suggest that there is widespread error and cryptic site usage by the mammalian splicing machinery, with unexplored roles for stochastic biological noise in shaping human transcriptome diversity. Previous measurements from steady-state RNA sequencing data have estimated that an average of ∼2% of transcription products from a gene might be erroneously spliced.^23^ Our results indicate that this is likely an underestimate and that it is necessary to profile splicing intermediates before they have a chance to be degraded by either nuclear or cytoplasmic quality control mechanisms to obtain a more complete picture of cryptic splicing products.^3^ By doing so, we are able to characterize the genomic features associated with 3 different categories of cryptic splicing: sites with high-fidelity, recursive, or low-fidelity spliceosome binding motifs.

This work challenges the widespread perspective that all alternative RNA splicing events or isoforms must be regulated and potentially have biological function. With the advent of very high-depth RNA-seq approaches, these data have been used to identify novel splicing events with high-fidelity spliceosome motifs but relatively low expression. Our results support the hypothesis put forward by previous modeling studies that the majority of these alternative isoforms arise from pairing between canonical splice sites likely caused by stochastic splicing decisions.^12,57^ Any significant fluctuations in the levels of these products might be consequences of direct regulation of the major isoform, fluctuations in the factors regulating splice site usage, or degradation of unproductive isoforms. Consistent with a recent study that used nascent RNA-seq to study unproductive splicing in human cells,^29^ we find that cryptic transcripts with high-fidelity splice sites appear to be more stable through the RNA lifecycle and are likely targeted by translation-dependent degradation processes (*i.e.* NMD) (**Figure 5D**). A subset of these sites, marked by recursive splice site motifs, might result in multi-step splicing of an intron. However, these putative recursive sites appear to share most of the characteristics of other high-fidelity cryptic sites. This is in line with recent results that despite the presence and usage of many sites with the potential for recursive splicing, bona fide recursive splicing happens at a very low-frequency in human cells^26^ and suggests that usage of these sites more often results in unstable, unproductive transcripts.

The majority of cryptic sites that we identify in newly created RNA use low-fidelity spliceosome motifs. These events must somehow bypass early kinetic proofreading steps, suggesting that there is heterogeneity in the initial spell-checking steps that evaluate surviving transcripts. This is consistent with the idea that kinetic proofreading cannot be perfect, at the risk of also reducing the production rate of productive transcripts if selection criteria are too stringent. Instead, there is likely a balance between productive and unproductive transcripts that survive proofreading steps. The existence of these cryptic sites points to an unappreciated role for stochasticity in the sequence recognition or molecular interactions that underlie splicing mechanisms.^23^ However, it remains unclear if there are genetic or biochemical factors that promote the usage of low-fidelity splice sites. These sites may be more representative of motifs recognized by variant or modified snRNA, which could have variable sequence preferences relative to canonical snRNAs.^58–60^ Given that we observe specific dinucleotide pairs that are more often used as low-fidelity splice sites (**Figure 3D,E**), it is possible that local sequence composition contributes to the usage of these sites, perhaps by promoting favorable structural conformations of the spliceosome to allow intron excision. Alternatively, it is also possible that local sequence composition influences RNA secondary structures — such as G-quadruplexes^38^ or base-pairing interactions between flanking regions — that create splicing-competent conformations.

Our results suggest that transcripts with low-fidelity splicing are more likely to be turned over quickly in every cellular compartment, especially in the nucleus (**Figure 5B,C**). Given that the splicing motifs themselves are spliced out of final transcripts, the question remains: what leads to transcripts that use different cryptic sites to be flagged for degradation with variable spatio-temporal dynamics? One possibility is there are changes in or a lack of EJC deposition at exon-exon junctions mediated by low-fidelity splice sites. This could involve differential EJC composition at these sites, consistent with our observation that knockdown of known EJC components (*i.e.* EIF4A3) causes an increase in the abundance of overall cryptic transcripts (**Figure 5A**). The sequence landscape around a junction might also influence EJC binding. A recent study found that m6A levels were depleted around canonical splice sites in pre-mRNA in a manner that enabled EJC deposition.^61^ Consequently, m6A modification near cryptic sites may prevent EJC recruitment or deposition. In a similar vein, we see a depletion of low-fidelity sites in a spliceosome- or EJC-sized window around regulated splice sites (**Figure 2D**). More broadly, it is unclear precisely how transcripts with low-fidelity sites are degraded. Since the RNA exosome is known to be involved in nuclear degradation of erroneous RNA products,^19^ this is the most obvious candidate for degradation of cryptic transcripts in the nucleus. However, further experiments are needed to test and confirm this hypothesis.

Finally, cryptic splicing seems to be linked to increased genomic real estate devoted to non-coding regions. This supports the multilevel selection theory that the prevalence of noisy splicing might be tolerated as a playground for evolutionary innovation even if only a small number of cryptic sites result in fitness advantages.^27,62^ Thus, some waste of metabolic resources to create unproductive transcripts might not be onerous depending on the function of the gene product or cellular context. This is consistent with our observation that genes involved in core cellular functions (*i.e.* cytoplasmic translation) tolerate less cryptic splicing, while lncRNA and expressed pseudogenes have less precise splicing regulation (**Figure 4C; Figure S6D**). It is unclear, however, whether these constraints occur at the level of splice site selection, quality control, or degradation and how specific genes are differentially regulated. Similarly, it is possible that cryptic splicing might be more advantageous in other cell types. We see cell-type specific variability in the abundance of cryptic site usage across genes — even for low-fidelity sites — suggesting that certain cellular environments might be more or less permissive of splicing noise. Cryptic splicing might be even more prevalent in cellular contexts or diseases such as cancer, where transcriptional noise could tax the splicing machinery and result in more chances for splicing error.^63,64^ Our results have established substantial variability in cryptic site usage across genes and cells, but more work is necessary to understand the causes, consequences, and landscape of cryptic splicing in mammalian cells.

## LIMITATIONS OF THE STUDY

To identify cryptic splice sites, we developed a computational framework called *CRYPTID-SS*. Within this workflow, we tried to account for (and remove) many artefactual sources of cryptic junction reads — including transposon or sequence variability, biases created during RNA handling or library preparation and sequencing or mapping errors — using control datasets to benchmark the probability that these issues may arise. It is possible that other unknown biological or technical causes of spurious junction reads may exist, which may affect the confidence in the positioning or quantification of any given lowly expressed cryptic site. However, any such biases should be consistent across genes and/or RNA enrichment types and have minimal effects on our results from analyses of aggregate genomic patterns.

## METHODS

### Human cell culture

K562 cells were grown in IMDM supplemented with 10% FBS. KNS60 cells were grown in DMEM supplemented with 5% FBS. AM38 cells were grown in EMEM supplemented with 20% FBS. HepG2 cells were grown in DMEM supplemented with 10% FBS. SH-SY5Y cells were grown in DMEM:F12 supplemented with 10% FBS. All cell lines were maintained at 37°C and 5% CO_2_ until 70% confluent. For cychoheximide experiments, K562 or KNS60 cells were treated 50ug/mL cycloheximide (CHX) (Sigma, C7698) resuspended in DMSO for 3 hours (or only DMSO as a control). For NMDi14 experiments, K562 and KNS60 cells were treated with 50uM NMDi14 (Sigma, SML1538) resuspended in DMSO for 6 hours. For transcriptional noise experiments, K562 cells were treated with 10uM 5-Iodo-2’-deoxyuridine (IdU) resuspended in DMSO for 24 hours and media containing 1mM 4sU (Sigma, T4509) was added during the last 8 minutes for subsequent nuclear isolation and nascent mRNA enrichment, as described below. Control cells were treated with equivalent amounts of DMSO without CHX, NMDi14, or IdU for 3, 6, and 24 hours, respectively.

### Nuclear isolation

Nuclear isolation from total cells was performed following a previously established protocol.^65^ Briefly, trypsinized cells were pelleted, washed with ice cold 1X PBS, and resuspended in ice cold Nuclear Isolation Buffer (10mM Tris-HCl, pH 7.4, 10mM NaCl, 3mM MgCl_2_, 0.3% NP40, Roche EDTA-free protease inhibitor (Millipore Sigma, 11836153001), Sigma PhosSTOP phosphatase inhibitor (Millipore Sigma, 4906845001). The suspension was briefly centrifuged again to pellet the nuclei. The supernatant was removed (cytoplasmic fraction) and the pellet was resuspended in ice cold Nuclear Isolation Buffer. The suspension was centrifuged at 10,000 RCF and supernatant removed. The nuclear pellet was resuspended in TRIzol (Invitrogen, 15596018) before RNA isolation.

The purity of the nuclear isolation was confirmed using a Western blot. To do so, cells were washed with ice-cold PBS and resuspended in ice-cold RIPA buffer containing EDTA-free protease cocktail inhibitor. Cell membranes were ruptured by sonication and clarified by centrifugation at 14,000 x g for 15 mins at 4°C. 10-20 µg cell lysate was loaded in each well and resolved on a 4-12% Tris-Glycine SDS PAGE and transferred to a nitrocellulose membrane. Briefly, the membrane was blocked in 5% milk, probed by antibodies (Cell Signaling Technologies, CST) 1:1,000 anti-vinculin and 1:1,000 anti-β-actin, followed by 1:2,000 anti-rabbit secondary (CST). HRP-linked secondary antibodies were visualized using ECL substrate (ThermoFisher).

### RNA isolation and library preparation

For nascent RNA isolation from nuclei, K562 and KNS60 cells were incubated with 500uM or 1mM 4-thiouridine (Sigma, T4509), AM-38, HepG2 and SY5Y cells were incubated in 1mM 4-thiouridine, dissolved in water, for 8 minutes and spun down to pellet cells. The pellet was resuspended in Trizol and flash frozen in liquid nitrogen. After nuclear isolation and RNA isolation and described below, 4sU pulldowns were performed using MTSEA biotin-XX (VWR, 89139-636) 4sU-containing (nascent) transcripts were purified following previously established protocols.^66,67^ Control cells were treated 1mM uridine (ChemGenes, RP-1186) dissolved in water for 8 minutes or with 1mM 4sU for 24 hours.

RNA was extracted from cells or nuclear pellets using TRIzol reagent to separate the aqueous RNA phase from the DNA and protein phases. The aqueous RNA phase was carried forward into column purification using a Zymo-25 RNA Clean & Concentrator kit (Zymo, R1018). Library preparation for mature RNA, CHX-treated, NMDi14-treated, and DMSO-treated samples was performed using the KAPA Stranded mRNA-Seq Kit for Illumina Platforms (KK8421). Library preparation for the nuclear and 4sU-labeled samples was performed using Kapa Biosystems KAPA RNA HyperPrep Kit with RiboErase (KK8561). Libraries were sequenced commercially at Novogene using an Illumina Novaseq or on an Illumina Nextseq 2000.

### Pre-processing and quantification of reads

Adapters were trimmed and removed from reads with trimmomatic v0.32. Adapter removal was verified by running fastqc v0.11.5^68^ on trimmed fastqc files. Overlapping read-mates were merged with PEAR v0.9.11^69^ and merged overlapping read-mates, non-overlapping read 1 and non-overlapping read 2 for each sample were pooled into a single fastq file per library. For all subsequent analyses, fastq files for cell lines were subsampled in two ways using seqtk v1.3^70^ and then processed as described above: (1) (subsampling to 112.67M reads to match the depth of the library with the lowest sequencing depth and (2) subsampling to percentiles of the total number of reads 10%, 20%, 30%, 40%, 50%, 60%, 70%, 80% and 90%) to analyze the sensitivity of cryptic site detection. Fastq files for K562 transcriptional noise experiments were subsampled to 33.1M reads to match the depth of library with the lowest sequencing depth.

Fastq files for SIRV synthetic RNA were not subsampled. Transcripts per million (TPM) values were calculated using kallisto v0.45.0^71^ on PEAR-processed, subsampled fastq files. Gene counts were generated from transcript counts from kallisto by tximport using EnsDB.Hsapiens.v86 annotations. Subsampled reads were subsequently processed using the CRYPTID-SS pipeline as described below. The distribution of intronic reads was estimated using bedtools coverage v2.28.0^72^ (with -s parameter) for all introns expressed genes (TPM >= 2).

### Identification of cryptic splice sites using *CRYPTID-SS*

We developed a python-based framework to confidently identify cryptic splice sites from RNA-seq exon-exon junction reads. Briefly, this pipeline does the following five steps, each of which are detailed below as applied to the datasets analyzed in this manuscript: (1) modified read names in fastq files to facilitate downstream analyses, (2) map reads with STAR splicing-aware mapper to identify and quantify junction reads, (3) resolve strand discrepancies for non-strand-specific junctions, (4) identify gene-specific and genomic features associated with each junction, (5) filter out junctions which are likely the result of biological or technical artefacts. *CRYPTID-SS* takes raw fastq files as an input and outputs a filtered table that contains junction, splice site, and genomic features.

<u>Step 1: Modifying read names.</u> By default, STAR trims the second part of read names using a space delimiter. As a result, the read orientation information (read 1 vs. read 2) is not carried forward to BAM files. To retain this information to aid in strand assignment (below), the space character in read names was replaced by an underscore character in FASTQ files pre-mapping.

<u>Step 2: Mapping reads.</u> Reads with modified read names were mapped to the GRCH38 genome using STAR v2.7.0e^73^ in single-end mode using default parameters with the following modifications or additions: *--outSAMattributes All --outSJfilterOverhangMin 40 12 12 12 --outSJfilterCountUniqueMin 1 1 1 1 --alignIntronMin 100 --alignIntronMax 250000 --limitOutSJcollapsed 2000000 --outSJfilterDistToOtherSJmin 20 0 10 10 --alignSJstitchMismatchNmax 2 2 2 2.* Reads are mapped in single-end mode since PEAR processing removes pairs for most reads. For all downstream steps, the resulting STAR junction database (.sj.out.tab) file was used.

<u>Step 3: Resolve strand discrepancies.</u> The STAR junction database does not contain strand information for junctions derived from sites with non-canonical intron motifs. To assign strand for these splice junctions, uniquely mapping junction reads were extracted for each site with a non-canonical intron motif. For each site, the number of uniquely mapped reads mapping to plus and minus strands were counted and the strand with the greater number of reads was assigned as the sense strand for that junction site.

<u>Step 4: Identify gene-specific and genomic features associated with junctions.</u> For each splice site within a junction, the following information was obtained by comparing to ENSEMBL GRCH38.95 annotations:

- Presence in annotations
- Location within a gene; ENSEMBL ID and strand if within a gene
- If within a gene, location within a annotated intron or exon including alternative or constitutive status of feature
- If within an exon, location within annotated UTRs, coding regions (CDS), or terminal codon
- Distance of splice site to nearest upstream 5’ss, nearest upstream 3’ss, nearest downstream 5’ss and nearest downstream 3’ss

To obtain sequence information, +/-25 nt of DNA sequence around the splice site and the relevant dinucleotide sequences (first 2nt downstream of 5’ sites or first 2nt upstream of 3’ sites) were extracted from the reference genome using *bedtools getfasta*. To estimate splice site strength, maxEnt scores for 5’ss (9nt region) and 3’ss (23nt region) were calculated using maxEntScan perl scripts that are modified to prevent skipping of sequences with “Ns” in them.^74^ To estimate conservation, base-specific and mean phyloP scores were obtained for a +/- 20nt region around the splice site by applying the *bigWigToBedGraph* tool from *ucsctools* using the hg38.phyloP100way bigwig file downloaded from the UCSC database.^75,76^

<u>Step 5: Filtering our spurious junctions.</u> The following criteria were applied to remove spruious splice junctions before downstream analyses:

- Splice junctions with sites within annotated intergenic regions.
- Splice junctions whose 5′ and 3′ splice sites mapped to different genes.
- Splice junctions with 5′ and 3′ splice sites mapping to a distance of less than 25 nucleotides.
- Splice junctions with no uniquely mapped reads.
- Splice junctions whose splice sites map within and near (within 10nt) the same repeat regions.
- Splice junctions whose 5′ and 3′ splice sites map within the same type of repeat within the same gene.
- Splice junctions containing polymorphisms in the first two or last two nucleotides of the intron (see genotyping section below).
- For K562 datasets, splice junctions containing indivdual splice sites that were also identified in the K562 exome dataset were removed.

### Identifying splice junctions within or near repeats

Bed file containing DNA repetitive elements for GRCh38 was downloaded from UCSC table browser. Splice junctions completely within repeat regions were identified with GenomicRanges using *findOverlaps( type = "within"*). Splice junctions near repeats with identified with GenomicRanges *findOverlaps( type = "within", maxgap=10*).

### Genotyping of cell lines from RNA-seq data

To ensure compatibility with GATK tools,^77^ MAPQ scores in BAM files generated from STAR were changed to “60” from “255”, and readgroups were added by *AddOrReplaceReadGroups* (picard v2.27.5).^78^ Junction reads were split using *SplitNCigarReads* (GATK v4.3.0.0).^79^ Haplotypes were called by *HaplotypeCaller* with the flag “*-ERC GVCF*” (GATK v4.3.0.0) and GVCF files from nuclear, nascent and polyA samples for each cell type (HepG2, K562, KNS60, AM38 and SY5Y) were combined using *CombineGVCFs* (GATK v4.3.0.0). Variants were cataloged using *GenotypeGVCFs* (GATK v4.3.0.0). Finally, variant .vcf files were converted to table format with *VariantstoTable* (GATK v4.3.0.0).

### Defining cryptic splice sites

Splicing junctions identified by *CRYPTID-SS* were collapsed into a unique set of splice sites, retaining junction pairing information for downstream analyses and position within the junction (5’ or 3’ site). Cryptic sites were categorized as belonging to one of three subtypes. High-fidelity cryptic junctions were defined as splice sites with canonical dinucleotides but have not been previously annotated as splice sites. These sites are derived from reads with SAM flag attributes of 𝑗𝑀: 𝐵: 𝑐, [1, 2, 3, 4, 5, 6]. Recursive sites were defined as those falling within an “AGGT” sequence motif necessary for recursive splicing, with 5’ recursive sites defining an intron beginning with the “GT” and 3’ recursive sites defining an intron ending with the “AG”. These sites are derived from reads with SAM flag attributes of 𝑗𝑀: 𝐵: 𝑐, [1, 2] and additionally have a templated 3’ or 5’ dinucleotide upstream of downstream of the 5’ or 3’ site, respectively. Low-fidelity cryptic sites were defined as splice sites without canonical dinucleotides (*i.e.* the first two nucleotides of the intron are not “GT” or “GC” or “AT” (for 5’ sites) and last two nucleotides of intron not containing “AG” or “AC” (for 3’ sites)). These sites are derived from reads with SAM flag attributes of 𝑗𝑀: 𝐵: 𝑐, [0, 20].

### Synthetic control data generation and analysis

Synthetic Spike-In RNA Variant (SIRV-Set1; Lexogen 025.03 & SIRV-Set4: Lexogen 141.01) was used to make libraries using both the stranded mRNA-seq and RNA HyperPrep kits, which were sequenced on NextSeq500. Resulting data was pre-processed and mapped as described above, using the SIRV reference annotations, and *CRYPTID-SS* was used to identify cryptic sites. SIRV data was used to estimate a false discovery rate (FDR) for errors introduced during library preparation and sequencing. Specifically, the FDR was calculated as the proportion of the false splicing instances (number of junction reads mapping to cryptic sites) among all splicing instances (total number of junction reads).

### Modeling major contributors to variability in usage of individual splice sites

To estimate the extent to which genomic factors at or around each splice site accounted for variance in the usage of splice sites, we fit a linear model of the following form, separately for canonical, low-fidelity, recursive, and high-fidelity splice sites:

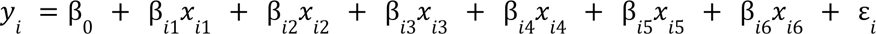

Where y_i_ reflects the usage of a splice site (log10, reads), x_i1_ is conservation score (mean phyloP score in a 50nt region around the splice site), x_i2_ is splice site score (from maxEnt), x_i3_ is the length of the intron or exon within which the site is in (log10, nucleotides), x_i4_ is the number of partner sites detected, x_i5_ is the mean distance between splice site partners (log10, nucleotides), and x_i6_ is the fractional position of the sites within the parent gene. We used the values from this multiple linear regression model to estimate the relative importance of each parameter contributing to variance in splice site usage. To do so, we used the *relaimpo* package in the R statistical environment,^80^ which arrives at a relative importance percentage by averaging the sequencing sum-of-squares obtained from all possible orderings of the predictors in the model.

### Co-transcriptional splicing analysis

The extent of co-transcriptional splicing that occurs at cryptic splice sites was estimated with individual indices for introns and exons. The completed splicing index (coSI)^81^ was calculated with read counts that span across exon boundaries into the adjacent intron (incomplete splicing) relative to read counts of split reads across exon-exon junctions (complete splicing). The coSI was used to assess co-transcriptional splicing when cryptic sites occurred in an intronic region. The splicing per intron (SPI)^82^ was calculated with junction read counts for spanning an exon-exon junction (completed splicing) relative to the total number of reads spanning exon-exon and exon-intron junctions (incompleted splicing). The SPI was used to assess co-transcriptional splicing when cryptic sites occurred in an exonic region.

### Dinucleotide analyses

All donor and acceptor dinucleotide pairs were extracted from the reference genome (either human or the SIRV reference sequence) and then randomly shuffled. The dinucleotides were selected at random. This shuffling process was repeated 1000 times to calculate a probability for each possible permutation in order to generate a null distribution. This null distribution was then used to calculate the statistical probability for the occurrence of dinucleotide pairs and identify significantly enriched dinucleotide pairings in the observed data.

### Counting the number of transcripts per cryptic splicing type

To estimate how many transcripts for each gene had type of cryptic splicing, we calculated TPMs using only junction reads, separated by cryptic type. We assumed that each transcript has at most one cryptic splicing event, and thus cryptic junction reads were treated as an individual transcript. Enrichment method-specific RPKMs were then calculated for each splice site category individually, where R is junction reads, K is the effective transcript length in kilobases estimated by *kallisto*, and M is the total number of junction reads (in millions). RPKMs for each gene were then summed up per sample to calculate junction-specific transcripts per million (TPM_junction_) as follows: 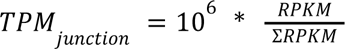.

### Elastic net regression modeling for genes

To build a predictive model for the extent of cryptic splicing per gene, we trained an elastic net regression model using the *caret* package in the R statistical environment.^83^ Briefly, we used the *glmnet* method to tune both the λ and α parameters using five-fold cross-validation and data were centered and scaled prior to use in the modeling. The variables modeled were transcription levels (log10, TPMs from nascent RNA), steady-state gene expression levels (log10, TPMs from mature RNA), gene length (log10, nucleotides), the number of introns in the gene, and the percentage of a gene that is intronic. Elastic net regression was chosen to select predictive features and avoid overfitting with variables that are highly correlated to each other. Expected values for each gene were predicted from the final model using the *predict* function in the R statistical environment and R^2^ values were calculated as the Pearson correlation between the predicted and observed values for each gene. Models were fit separately to data from each enrichment method and each category of splice sites (canonical, low-fidelity, recursive, and high-fidelity sites). To identify genes with more cryptic splicing than expected given the variables modeled, studentized residuals were calculated from the model residuals and any gene with a studentized residual greater than 1.5 was categorized as having more cryptic splicing than expected for that cryptic splicing category.

### Gene Ontology

Gene ontology analysis was performed using WEB-based GEne SeT AnaLysis Toolkit^84^ to perform gene set enrichment analysis for genes in the categories defined by junction TPMs and predicted outliers characterized by the elastic net regression model.

### Analysis of ENCODE RBP data

FASTQ files for shRNA knockdown RNA-seq samples against RBPs in K562 cells were downloaded from the ENCODE portal.^43^ FASTQ files from read1 and read2 were processed with PEAR, subsampled to 12.7M reads, and mapped with STAR as described above. Cryptic splice sites were identified with CRYPTID-SS and filtered as described above. The proportions of cryptic splice sites and cryptic junction reads in each knockdown were compared with the batch-matched scrambled shRNA control sample. To determine statistical significance, control shRNA samples were compared to other randomly selected control shRNA samples 100 times to produce null distributions of the proportions of cryptic sites and cryptic junction reads. The observed proportion of cryptic sites or reads in the RBP KD samples were compared to the null distribution to determine the probability of observing these proportions by chance. FDRs for p-values were calculated using the Benjamini Hochberg correction.

### Analysis of RNA flow data

To assess the lifecycle of genes with more cryptic sites than expected, we used subcellular residence times and half-lives estimated for each gene in K562 cells from the RNAflow method as found in Supplementary Table 1 from Ietswaart *et al.*^56^ Predicted to Undergo Nuclear Degradation (PUND) classification for each gene was also obtained from Ietswaart *et al.*^56^

## CODE & DATA AVAILABILITY

The CRYPTID-SS pipeline and code for downstream analyses included in this manuscript are available at https://github.com/thepailab/CRYPTID-SS. Sequencing data generated for this study are available at the Gene Expression Omnibus (GSE302375).

## ACKNOWLEDGEMENTS

We thank E. Sontheimer, P. Zamore, and Watts and Pai labs for helpful discussions and comments. This work was funded by grants from the National Institutes of Health (R35GM133762 A.A.P. and R01NS111990 to J.K.W.), the National Science Foundation (CAREER2237568 to A.A.P.), and the SLC6A1 Connect Foundation.

## AUTHOR CONTRIBUTIONS

Contributed equally (EK, KB), Conceived and designed experiments (JKW, AAP), Performed the experiments (KB, ZK, VS, NJ), Performed statistical analyses (EK, AK, AAP), Analyzed the data (EK, AK, AAP), Contributed reagents/materials/analysis tools (JKW, AAP), Wrote the paper (KB, AAP), Jointly supervised research (JKW, AAP).

## REFERENCES

1. Hollander, D., Naftelberg, S., Lev-Maor, G., Kornblihtt, A.R., and Ast, G. (2016). How Are Short Exons Flanked by Long Introns Defined and Committed to Splicing? Trends Genet. 32, 596–606.

2. Saudemont, B., Popa, A., Parmley, J.L., Rocher, V., Blugeon, C., Necsulea, A., Meyer, E., and Duret, L. (2017). The fitness cost of mis-splicing is the main determinant of alternative splicing patterns. Genome Biol. 18, 208.

3. Wan, Y., and Larson, D.R. (2018). Splicing heterogeneity: separating signal from noise. Genome Biol. 19, 86.

4. De Conti, L., Baralle, M., and Buratti, E. (2013). Exon and intron definition in pre-mRNA splicing. Wiley Interdiscip. Rev. RNA 4, 49–60.

5. Dvinge, H. (2018). Regulation of alternative mRNA splicing: old players and new perspectives. FEBS Lett. 592, 2987–3006.

6. Wang, Z., and Burge, C.B. (2008). Splicing regulation: from a parts list of regulatory elements to an integrated splicing code. RNA 14, 802–813.

7. Bentley, D.L. (2014). Coupling mRNA processing with transcription in time and space. Nat. Rev. Genet. 15, 163–175.

8. Carey, L.B. (2015). RNA polymerase errors cause splicing defects and can be regulated by differential expression of RNA polymerase subunits. 10.7554/eLife.09945.

9. Gout, J.-F., Li, W., Fritsch, C., Li, A., Haroon, S., Singh, L., Hua, D., Fazelinia, H., Smith, Z., Seeholzer, S., et al. (2017). The landscape of transcription errors in eukaryotic cells. Science Advances. 10.1126/sciadv.1701484.

10. Mohler, K., and Ibba, M. (2017). Translational fidelity and mistranslation in the cellular response to stress. Nature Microbiology 2, 1–9.

11. Fox-Walsh, K.L., and Hertel, K.J. (2009). Splice-site pairing is an intrinsically high fidelity process. Proceedings of the National Academy of Sciences 106, 1766–1771.

12. Melamud, E., and Moult, J. (2009). Stochastic noise in splicing machinery. Nucleic Acids Res. 37, 4873–4886.

13. Semlow, D.R., and Staley, J.P. (2012). Staying on message: ensuring fidelity in pre-mRNA splicing. Trends Biochem. Sci. 37, 263–273.

14. Skandalis, A. (2016). Estimation of the minimum mRNA splicing error rate in vertebrates. Mutat. Res. 784–785, 34–38.

15. Davidson, L., Kerr, A., and West, S. (2012). Co-transcriptional degradation of aberrant pre-mRNA by Xrn2. EMBO J. 31, 2566–2578.

16. Egecioglu, D.E., and Chanfreau, G. (2011). Proofreading and spellchecking: a two-tier strategy for pre-mRNA splicing quality control. RNA 17, 383–389.

17. Thomas, M.J., Platas, A.A., and Hawley, D.K. (1998). Transcriptional fidelity and proofreading by RNA polymerase II. Cell 93, 627–637.

18. Sydow, J.F., and Cramer, P. (2009). RNA polymerase fidelity and transcriptional proofreading. Curr. Opin. Struct. Biol. 19, 732–739.

19. Schmid, M., and Jensen, T.H. (2018). Controlling nuclear RNA levels. Nature Reviews Genetics 19, 518–529.

20. Morris, C., Cluet, D., and Ricci, E.P. (2021). Ribosome dynamics and mRNA turnover, a complex relationship under constant cellular scrutiny. Wiley Interdisciplinary Reviews: RNA 12, e1658.

21. Monaghan, L., Longman, D., and Cáceres, J.F. (2023). Translation-coupled mRNA quality control mechanisms. The EMBO Journal. 10.15252/embj.2023114378.

22. Dou, Y., Fox-Walsh, K.L., Baldi, P.F., and Hertel, K.J. (2006). Genomic splice-site analysis reveals frequent alternative splicing close to the dominant splice site. RNA 12, 2047–2056.

23. Pickrell, J.K., Pai, A.A., Gilad, Y., and Pritchard, J.K. (2010). Noisy Splicing Drives mRNA Isoform Diversity in Human Cells. PLoS Genet. 6, e1001236.

24. Duff, M.O., Olson, S., Wei, X., Garrett, S.C., Osman, A., Bolisetty, M., Plocik, A., Celniker, S.E., and Graveley, B.R. (2015). Genome-wide identification of zero nucleotide recursive splicing in Drosophila. Nature 521, 376–379.

25. Pai, A.A., Paggi, J.M., Yan, P., Adelman, K., and Burge, C.B. (2018). Numerous recursive sites contribute to accuracy of splicing in long introns in flies. PLoS Genet.

26. Wan, Y., Anastasakis, D.G., Rodriguez, J., Palangat, M., Gudla, P., Zaki, G., Tandon, M., Pegoraro, G., Chow, C.C., Hafner, M., et al. (2021). Dynamic imaging of nascent RNA reveals general principles of transcription dynamics and stochastic splice site selection. Cell 184, 2878–2895.e20.

27. Sibley, C.R., Blazquez, L., and Ule, J. (2016). Lessons from non-canonical splicing. Nat. Rev. Genet. 17, 407–421.

28. Zhang, Z., Xin, D., Wang, P., Zhou, L., Hu, L., Kong, X., and Hurst, L.D. (2009). Noisy splicing, more than expression regulation, explains why some exons are subject to nonsense-mediated mRNA decay. BMC Biol. 7, 23.

29. Fair, B., Buen Abad Najar, C.F., Zhao, J., Lozano, S., Reilly, A., Mossian, G., Staley, J.P., Wang, J., and Li, Y.I. (2024). Global impact of unproductive splicing on human gene expression. Nat. Genet. 56, 1851–1861.

30. Shenasa, H., and Bentley, D.L. (2023). Pre-mRNA splicing and its cotranscriptional connections. Trends Genet. 39, 672–685.

31. McManus, C.J., Duff, M.O., Eipper-Mains, J., and Graveley, B.R. (2010). Global analysis of trans-splicing in Drosophila. Proc. Natl. Acad. Sci. U. S. A. 107, 12975–12979.

32. Lim, K.H., Han, Z., Jeon, H.Y., Kach, J., Jing, E., Weyn-Vanhentenryck, S., Downs, M., Corrionero, A., Oh, R., Scharner, J., et al. (2020). Antisense oligonucleotide modulation of non-productive alternative splicing upregulates gene expression. Nature Communications 11, 1–13.

33. Bradnam, K.R., and Korf, I. (2008). Longer first introns are a general property of eukaryotic gene structure. PLoS One 3, e3093.

34. Sibley, C.R., Emmett, W., Blazquez, L., Faro, A., Haberman, N., Briese, M., Trabzuni, D., Ryten, M., Weale, M.E., Hardy, J., et al. (2015). Recursive splicing in long vertebrate genes. Nature 521, 371–375.

35. Zhang, X.-O., Fu, Y., Mou, H., Xue, W., and Weng, Z. (2018). The temporal landscape of recursive splicing during Pol II transcription elongation in human cells. PLoS Genet. 14, e1007579.

36. Hoppe, E.R., Udy, D.B., and Bradley, R.K. (2023). Recursive splicing discovery using lariats in total RNA sequencing. Life Sci Alliance 6. 10.26508/lsa.202201889.

37. Bradley, R.K., Merkin, J., Lambert, N.J., and Burge, C.B. (2012). Alternative splicing of RNA triplets is often regulated and accelerates proteome evolution. PLoS Biol 10, e1001229.

38. Georgakopoulos-Soares, I., Parada, G.E., Wong, H.Y., Medhi, R., Furlan, G., Munita, R., Miska, E.A., Kwok, C.K., and Hemberg, M. (2022). Alternative splicing modulation by G-quadruplexes. Nature Communications 13, 1–16.

39. Sheth, N., Roca, X., Hastings, M.L., Roeder, T., Krainer, A.R., and Sachidanandam, R. (2006). Comprehensive splice-site analysis using comparative genomics. Nucleic Acids Res. 34, 3955–3967.

40. Staněk, D. (2021). Long non-coding RNAs and splicing. Essays Biochem 65, 723–729.

41. Basu, K., Dey, A., and Kiran, M. (2023). Inefficient splicing of long non-coding RNAs is associated with higher transcript complexity in human and mouse. RNA Biology, 563–572.

42. Desai, R.V., Chen, X., Martin, B., Chaturvedi, S., Hwang, D.W., Li, W., Yu, C., Ding, S., Thomson, M., Singer, R.H., et al. (2021). A DNA repair pathway can regulate transcriptional noise to promote cell fate transitions. Science 373, eabc6506.

43. ENCODE Project Consortium (2012). An integrated encyclopedia of DNA elements in the human genome. Nature 489, 57–74.

44. Santonoceto, G., Jurkiewicz, A., and Szczesny, R.J. (2024). RNA degradation in human mitochondria: the journey is not finished. Hum Mol Genet 33, R26–R33.

45. Schlautmann, L.P., and Gehring, N.H. (2020). A Day in the Life of the Exon Junction Complex. Biomolecules 10, 866.

46. McCloskey, A., Taniguchi, I., Shinmyozu, K., and Ohno, M. (2012). hnRNP C tetramer measures RNA length to classify RNA polymerase II transcripts for export. Science 335, 1643–1646.

47. Liu, N., Dai, Q., Zheng, G., He, C., Parisien, M., and Pan, T. (2015). N6-methyladenosine-dependent RNA structural switches regulate RNA–protein interactions. Nature 518, 560–564.

48. Wang, X., Zhao, B.S., Roundtree, I.A., Lu, Z., Han, D., Ma, H., Weng, X., Chen, K., Shi, H., and He, C. (2015). N(6)-methyladenosine Modulates Messenger RNA Translation Efficiency. Cell 161, 1388–1399.

49. Hirose, T., Ideue, T., Nagai, M., Hagiwara, M., Shu, M.-D., and Steitz, J.A. (2006). A spliceosomal intron binding protein, IBP160, links position-dependent assembly of intron-encoded box C/D snoRNP to pre-mRNA splicing. Mol. Cell 23, 673–684.

50. De, I., Bessonov, S., Hofele, R., dos Santos, K., Will, C.L., Urlaub, H., Lührmann, R., and Pena, V. (2015). The RNA helicase Aquarius exhibits structural adaptations mediating its recruitment to spliceosomes. Nat. Struct. Mol. Biol. 22, 138–144.

51. Schmitzová, J., Cretu, C., Dienemann, C., Urlaub, H., and Pena, V. (2023). Structural basis of catalytic activation in human splicing. Nature 617, 842–850.

52. Beusch, I., and Madhani, H.D. (2024). Understanding the dynamic design of the spliceosome. Trends Biochem Sci 49, 583–595.

53. Splicing quality control mediated by DHX15 and its G-patch activator SUGP1 (2023). Cell Reports 42, 113223.

54. García-Ruiz, S., Zhang, D., Gustavsson, E.K., Rocamora-Perez, G., Grant-Peters, M., Fairbrother-Browne, A., Reynolds, R.H., Brenton, J.W., Gil-Martínez, A.L., Chen, Z., et al. (2023). Splicing accuracy varies across human introns, tissues and age. bioRxiv, 2023.03.29.534370. 10.1101/2023.03.29.534370.

55. Van Nostrand, E.L., Pratt, G.A., Yee, B.A., Wheeler, E.C., Blue, S.M., Mueller, J., Park, S.S., Garcia, K.E., Gelboin-Burkhart, C., Nguyen, T.B., et al. (2020). Principles of RNA processing from analysis of enhanced CLIP maps for 150 RNA binding proteins. Genome Biol 21, 90.

56. Ietswaart, R., Smalec, B.M., Xu, A., Choquet, K., McShane, E., Jowhar, Z.M., Guegler, C.K., Baxter-Koenigs, A.R., West, E.R., Fu, B.X.H., et al. (2024). Genome-wide quantification of RNA flow across subcellular compartments reveals determinants of the mammalian transcript life cycle. Mol. Cell 84, 2765–2784.e16.

57. Hu, J., Boritz, E., Wylie, W., and Douek, D.C. (2017). Stochastic principles governing alternative splicing of RNA. PLoS Comput. Biol. 13, e1005761.

58. Mabin, J.W., Lewis, P.W., Brow, D.A., and Dvinge, H. (2021). Human spliceosomal snRNA sequence variants generate variant spliceosomes. RNA 27, 1186–1203.

59. Parker, M.T., Fica, S.M., Barton, G.J., and Simpson, G.G. (2023). Inter-species association mapping links splice site evolution to METTL16 and SNRNP27K. 10.7554/eLife.91997.

60. Parker, M.T., Soanes, B.K., Kusakina, J., Larrieu, A., Knop, K., Joy, N., Breidenbach, F., Sherwood, A.V., Barton, G.J., Fica, S.M., et al. (2022). m6A modification of U6 snRNA modulates usage of two major classes of pre-mRNA 5’ splice site. 10.7554/eLife.78808.

61. Sethi, A.J., Guarnacci, M., Bilal, M., Krishnan, K.S., Hayashi, A., Kanchi, M., Nojima, T., Preiss, T., Eyras, E., and Hayashi, R. (2025). Single-molecule multimodal timing of*in vivo*mRNA synthesis. bioRxiv. 10.1101/2025.04.27.650906.

62. Brunet, T.D.P., and Doolittle, W.F. (2015). Multilevel Selection Theory and the Evolutionary Functions of Transposable Elements. Genome Biol. Evol. 7, 2445–2457.

63. Lin, C.Y., Lovén, J., Rahl, P.B., Paranal, R.M., Burge, C.B., Bradner, J.E., Lee, T.I., and Young, R.A. (2012). Transcriptional amplification in tumor cells with elevated c-Myc. Cell 151, 56–67.

64. Chen, L., Tovar-Corona, J.M., and Urrutia, A.O. (2011). Increased levels of noisy splicing in cancers, but not for oncogene-derived transcripts. Hum. Mol. Genet. 20, 4422–4429.

65. Nabbi, A., and Riabowol, K. (2015). Rapid isolation of nuclei from cells in vitro. Cold Spring Harb. Protoc. 2015, 769–772.

66. Garibaldi, A., Carranza, F., and Hertel, K.J. (2017). Isolation of newly transcribed RNA using the metabolic label 4-thiouridine. Methods Mol. Biol. 1648, 169–176.

67. Duffy, E.E., and Simon, M.D. (2016). Enriching s4 U-RNA using methane thiosulfonate (MTS) chemistry. Curr. Protoc. Chem. Biol. 8, 234–250.

68. Andrews, S. (2016). FastQC: A quality control tool for high throughput sequence data. https://www.bioinformatics.babraham.ac.uk/projects/fastqc/.

69. Zhang, J., Kobert, K., Flouri, T., and Stamatakis, A. (2014). PEAR: a fast and accurate Illumina Paired-End reAd mergeR. Bioinformatics 30, 614–620.

70. Li, H. (2018). seqtk. seqtk: Toolkit for processing sequences in FASTA/Q formats. https://github.com/lh3/seqtk.

71. Bray, N.L., Pimentel, H., Melsted, P., and Pachter, L. (2016). Near-optimal probabilistic RNA-seq quantification. Nat. Biotechnol. 34, 525–527.

72. Quinlan, A.R., and Hall, I.M. (2010). BEDTools: a flexible suite of utilities for comparing genomic features. Bioinformatics 26, 841–842.

73. Dobin, A., Davis, C.A., Schlesinger, F., Drenkow, J., Zaleski, C., Jha, S., Batut, P., Chaisson, M., and Gingeras, T.R. (2013). STAR: ultrafast universal RNA-seq aligner. Bioinformatics 29, 15–21.

74. Yeo, G., and Burge, C.B. (2004). Maximum entropy modeling of short sequence motifs with applications to RNA splicing signals. J. Comput. Biol. 11, 377–394.

75. Siepel, A., Bejerano, G., Pedersen, J.S., Hinrichs, A.S., Hou, M., Rosenbloom, K., Clawson, H., Spieth, J., Hillier, L.W., Richards, S., et al. (2005). Evolutionarily conserved elements in vertebrate, insect, worm, and yeast genomes. Genome Res. 15, 1034–1050.

76. Raney, B.J., Barber, G.P., Benet-Pagès, A., Casper, J., Clawson, H., Cline, M.S., Diekhans, M., Fischer, C., Navarro Gonzalez, J., Hickey, G., et al. (2023). The UCSC Genome Browser database: 2024 update. Nucleic Acids Res. 10.1093/nar/gkad987.

77. Van der Auwera, G.A., Carneiro, M.O., Hartl, C., Poplin, R., Del Angel, G., Levy-Moonshine, A., Jordan, T., Shakir, K., Roazen, D., Thibault, J., et al. (2013). From FastQ data to high confidence variant calls: the Genome Analysis Toolkit best practices pipeline. Curr. Protoc. Bioinformatics 43, 11.10.1–11.10.33.

78. Picard toolkit (2019). Broad Institute, GitHub repository.

79. Poplin, R., Ruano-Rubio, V., DePristo, M.A., Fennell, T.J., Carneiro, M.O., Van der Auwera, G.A., Kling, D.E., Gauthier, L.D., Levy-Moonshine, A., Roazen, D., et al. (2017). Scaling accurate genetic variant discovery to tens of thousands of samples. bioRxiv. 10.1101/201178.

80. Groemping, U. (2006). Relative importance for linear regression in R: The package relaimpo. Journal of Statistical Software 17, 1–27.

81. Tilgner, H., Knowles, D.G., Johnson, R., Davis, C.A., Chakrabortty, S., Djebali, S., Curado, J., Snyder, M., Gingeras, T.R., and Guigó, R. (2012). Deep sequencing of subcellular RNA fractions shows splicing to be predominantly co-transcriptional in the human genome but inefficient for lncRNAs. Genome Res. 22, 1616–1625.

82. Herzel, L., and Neugebauer, K.M. (2015). Quantification of co-transcriptional splicing from RNA-Seq data. Methods 85, 36–43.

83. Kuhn, M. (2008). Building Predictive Models inRUsing thecaretPackage. J. Stat. Softw. 28, 1–26.

84. Elizarraras, J.M., Liao, Y., Shi, Z., Zhu, Q., Pico, A.R., and Zhang, B. (2024). WebGestalt 2024: faster gene set analysis and new support for metabolomics and multi-omics. Nucleic Acids Res 52, W415–W421.

