## Supplementary Figures for "Pervasive noise in human splice site selection"

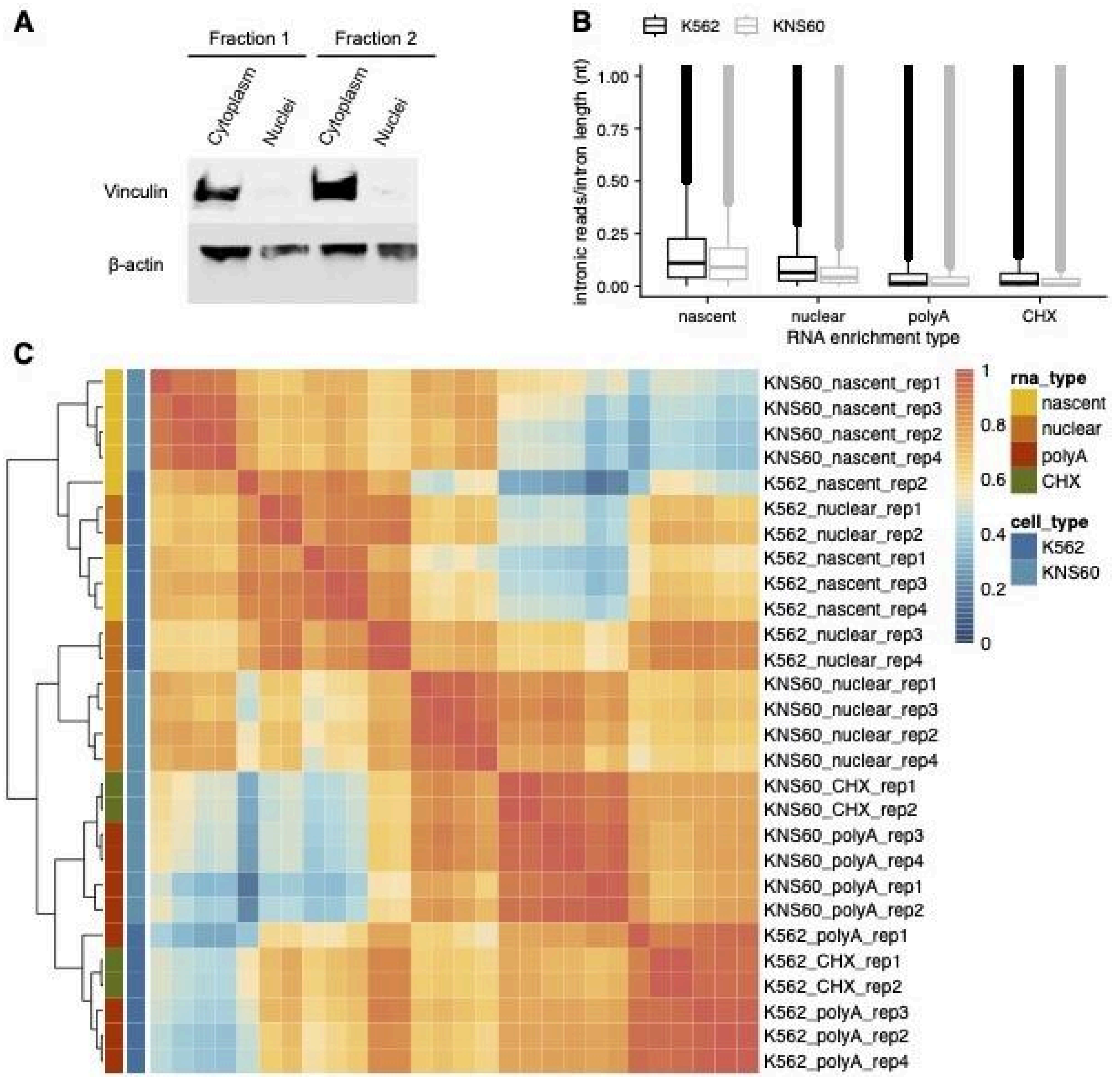


**Supplementary Figure 1. Validation of RNA enrichment methods. (A)** Western blot of fractionated cell samples. Vinculin, a membrane-cytoskeletal adhesion protein, is absent in purified nuclear fractions but present in cytoplasmic fractions. β-actin was used as a loading control. **(B)** Distribution of intronic reads (normalized to intron length, *y-axis)* across RNA enrichment methods *(x-axis)* for K562 *(black)* and KNS60 *(gray)* cells*.* **(C)** Clustering heatmap of TPM values for each sample across cell types and RNA enrichment methods, with a hierarchical clustering dendogram shown on the left*.*

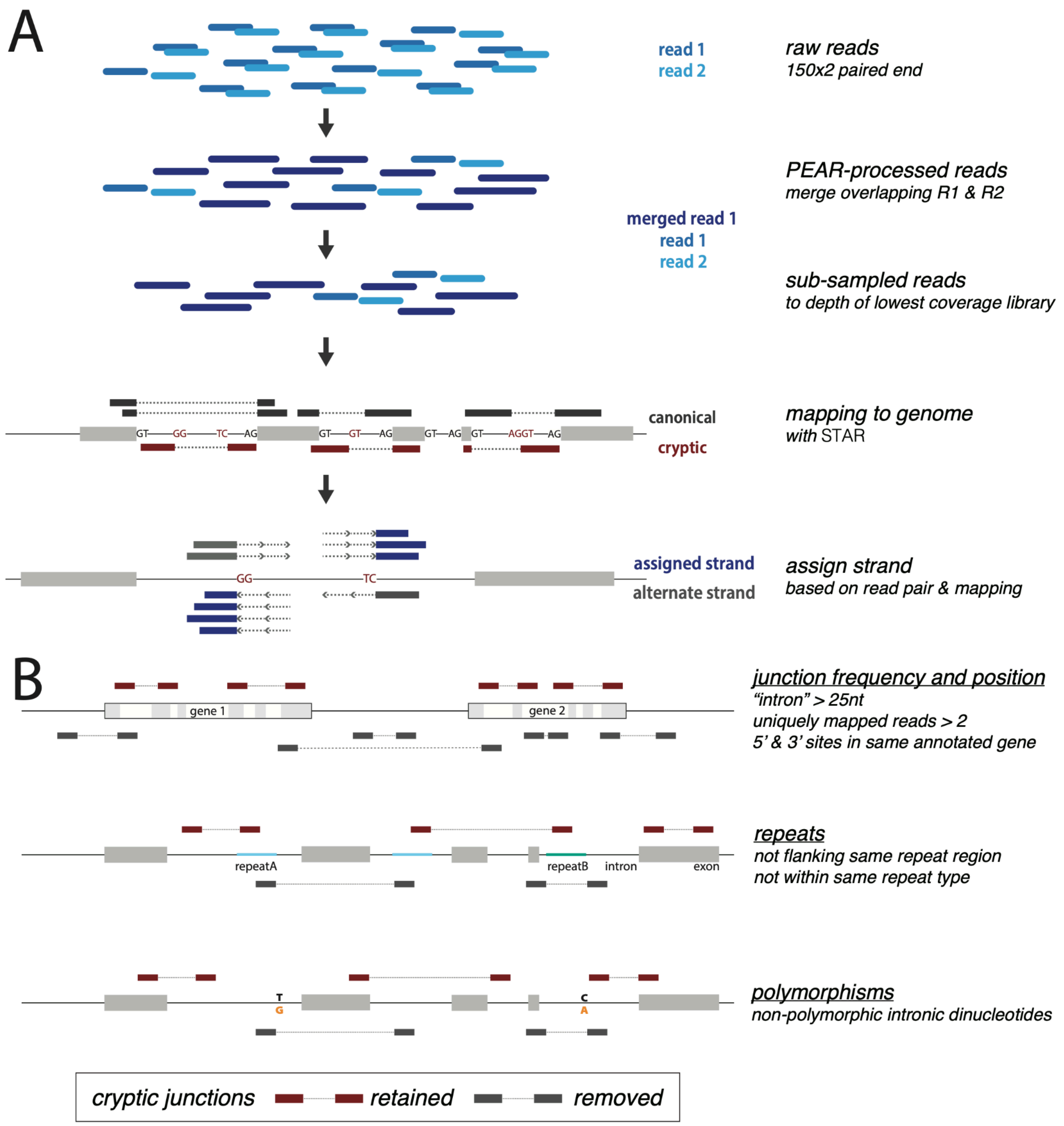


**Supplementary Figure 2. Computational pipelines and criteria used to identify cryptic splice sites. (A)** Schematic representation of the computational pipeline to identify cryptic splice sites, including collapsing overlapping read pairs, subsampling read depth, mapping and annotation, and strand assignment for novel sites. **(B)** Schematic representation of the criteria necessary to identify cryptic splice sites, including junction frequency and position, proximity to repeat region(s), and overlap with sequence variation.

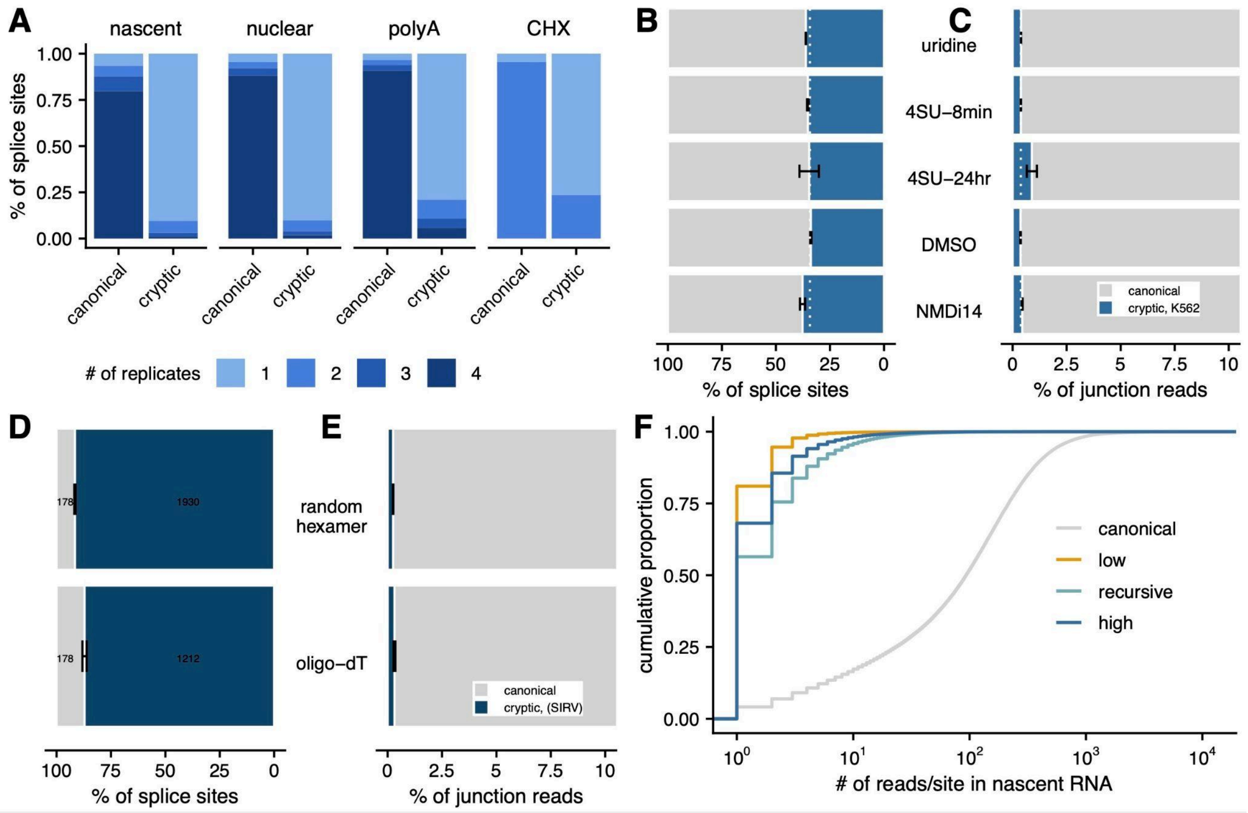


**Supplementary Figure 3. Control datasets used to establish confidence in cryptic site identification and quantification. (A)** Percentage of canonical or cryptic splice sites that were detected in 1, 2, 3, or all 4 replicates across enrichment methods. Percentage of canonical (*grey*) and cryptic (*blue*) splice sites **(B)** and junction reads **(C)** detected across control treatments in K562 cells. Dotted white line represents the mean percentage for polyA-enriched samples. Percentage of canonical (*grey*) and cryptic (*blue*) splice sites **(D)** and junction reads **(E)** detected in random hexamer or oligo-dT primed SIRV libraries. **(F)** Cumulative proportion (*y-axis*) of junction reads per site (*x-axis*) in nascent RNA for canonical *(gray)*, low-fidelity *(orange)*, recursive *(teal)*, and high-fidelity *(dark blue)* splice sites. **(G)** Percentage of canonical, low-fidelity, recursive, and high-fidelity cryptic splice sites that were detected in 1, 2, 3, or all 4 enrichment methods.

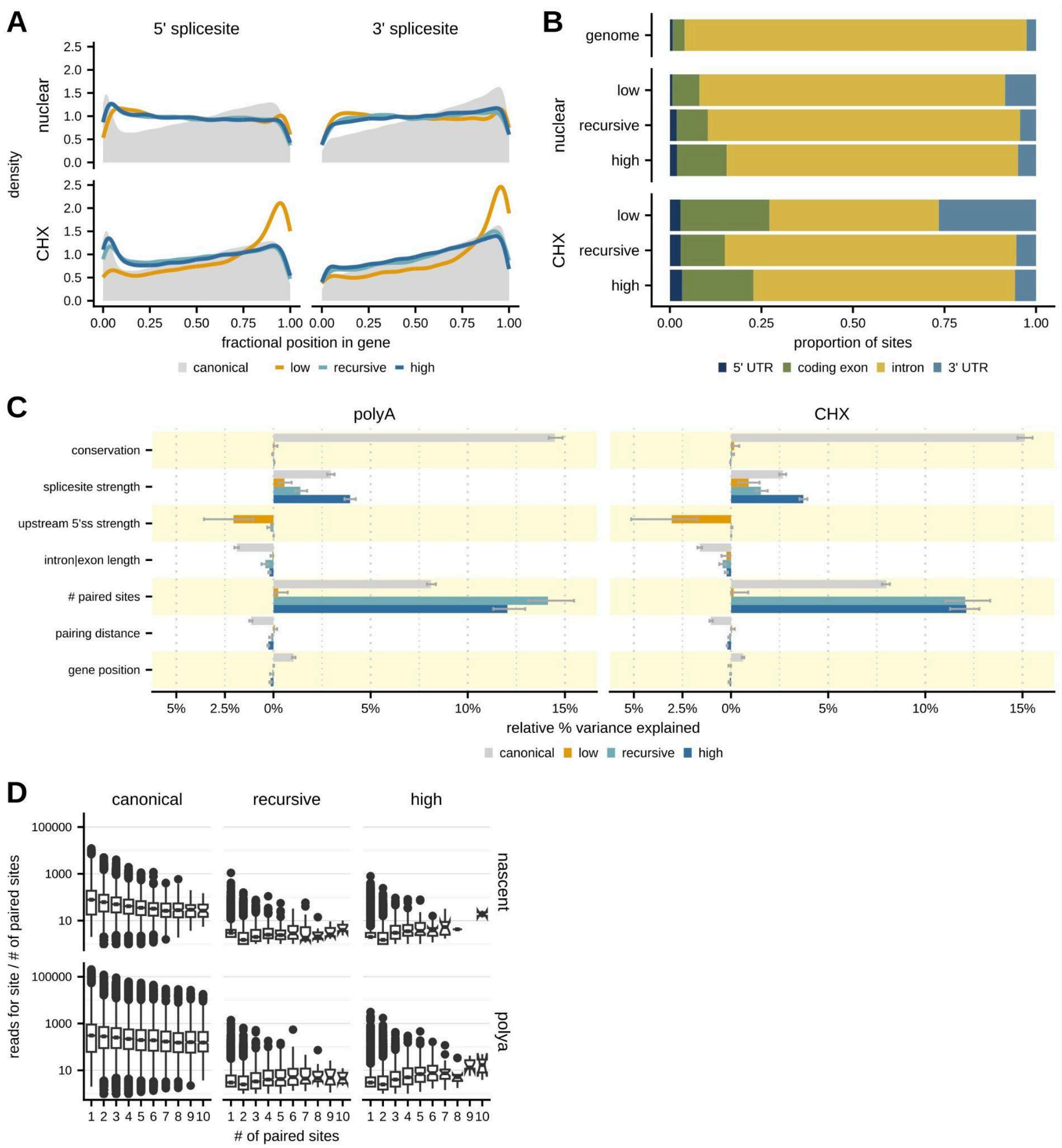


**Supplementary Figure 4. Genomic features of cryptic sites across RNA enrichment methods. (A)** Distribution of the position of 5' (*left*) and 3' (*right*) splice sites across a gene categorized as canonical (*grey*), low-fidelity (*orange*), recursive *(teal)*, or high-fidelity (*dark blue)* in either nuclear (*top*) or CHX-treated (*bottom*) RNA, where the fractional position is defined as the distance of a site from the start of a gene relative to the total gene length. **(B)** The proportion of each type of cryptic site detected within annotated 5' UTRs (*navy blue*), coding exons (*green*), introns (*yellow*), and 3' UTRs (*light blue*) for sites detected in nuclear (*middle*) or CHX-treated (*bottom*) RNA. The top bar shows the proportion of nucleotides within expressed genes for each annotation category. **(C)** Relative importance of features influencing variance in splice site usage for canonical (*grey*), low-fidelity (*orange*), recursive (*teal*), and high-fidelity (*dark blue*) sites, using multiple linear regression models for mature *(left)* and CHX-treated RNA *(right)*. Bars pointing to the right indicate positively correlated features, while bars pointing left indicate negatively correlated features. **(D)** Distribution of the number of reads relative to the number of paired sites for the same site *(y-axis)* across splice sites with increasing numbers of paired sites *(x-axis)* for canonical *(left)*, recursive *(middle)*, and high-fidelity *(right)* splice sites in nascent *(top)* and mature *(bottom)* RNA. Very few low-fidelity sites have more than one pairing partner.

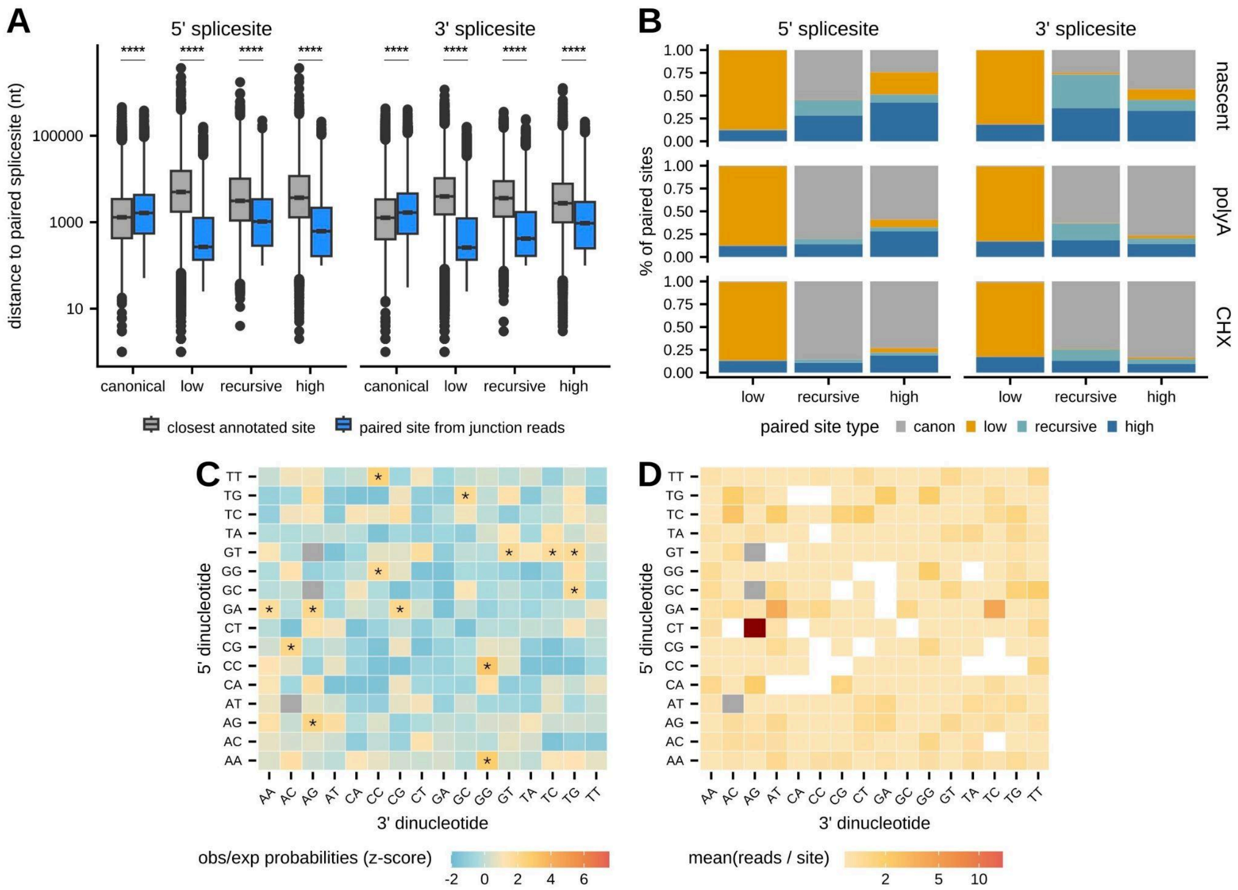


**Supplementary Figure 5. Pairing characteristics of canonical and cryptic splice sites. (A)** The distribution of distances between pairing partners (blue) or the closest annotated splice site (grey) for 5' *(left)* and 3' *(right)* splice sites divided by canonical, low-fidelity, recursive, and high-fidelity sites. Signficance was assessed with a Mann-Whitney U Test. **** adjusted p-value < 0.0001 **(B)** Proportion of 5' *(left)* and 3' *(right)* splice sites divided by cryptic type (*x-axis*) that pair with canonical *(gray)*, low-fidelity *(orange)*, recursive *(teal)*, or high-fidelity *(blue)* site in nascent *(top)*, mature *(middle)*, and CHX-treated *(bottom)* RNA. **(D)** Heatmap of the ratio between the observed number of sites relative to the expected distribution of sites (calculated by estimating the frequency each dinucleotide occurs in the SIRV transcriptome) for each combination of 5' (*y-axis*) and 3' (*x-axis*) dinucleotides used by non-annotated splice sites detected in SIRV libraries. Dinucleotide pairs involved in canonical splice site pairings were omitted from the plot (*grey boxes*). Signficance was assessed using permutations. * adjusted p-value < 0.05. **(E)** Heatmap of the mean number of reads per site for each combination of 5' (*y-axis*) and 3' (*x-axis*) dinucleotides used by non-annotated splice sites detected in SIRV libraries. Dinucleotide pairs involved in canonical splice site pairings and pairs with fewer than 25 observations were omitted from the plot (*grey* and *white boxes*, respectively).

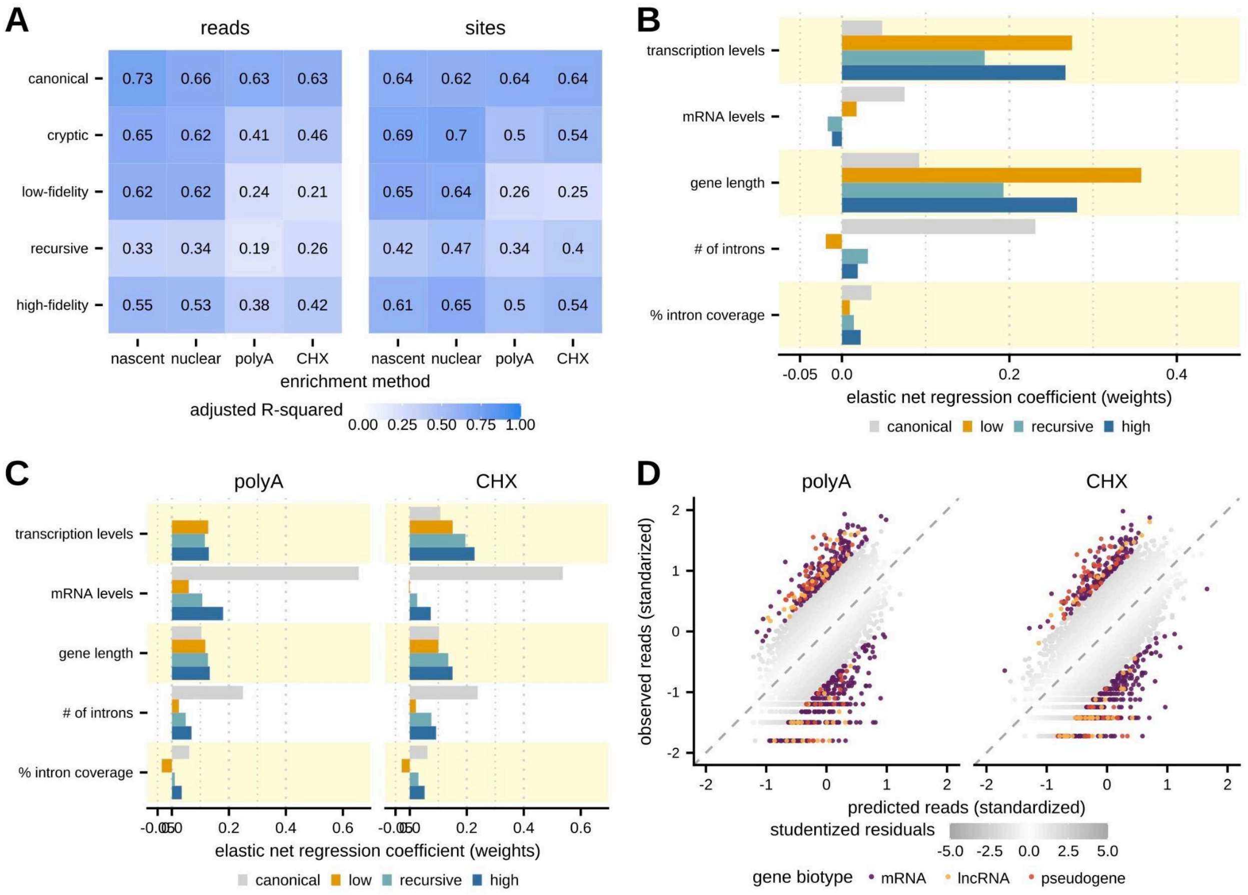


**Supplementary Figure 6. Elastic net regression to predict cryptic splicing across RNA enrichment methods. (A)** The adjusted R-squared values for elastic net regression models trained on different splice site (*y-axis*) in each enrichment method (*x-axis*) using the number of junction reads *(left)* or splice sites *(right).* **(B)** Feature weights from elastic net regression models (*x-axis*) trained on the number of cryptic splice sites per gene for canonical (*grey*), low-fidelity (*orange*), recursive (*teal*), and high-fidelity (*dark blue*) sites in nascent RNA. Features are described in Figure 4A. **(C)** Feature weights from elastic net regression models (*x-axis*) trained on the number of cryptic splicing reads per gene for canonical (*grey*), low-fidelity (*orange*), recursive (*teal*), and high-fidelity (*dark blue*) sites in polyA *(left)* and CHX-treated *(right)* RNA. Features are described in Figure 4A. **(D)** The amount of cryptic splicing per gene predicted by the elastic net regression model (center standardized, *x-axis*) versus the observed cryptic splicing per gene (*y-axis*) in polyA *(left)* and CHX-treated *(right)* RNA. The grey color scale indicates the magnitude of studentized residuals and genes with |studentized residual| ≥ 2 are colored by their ENSEMBL biotype.

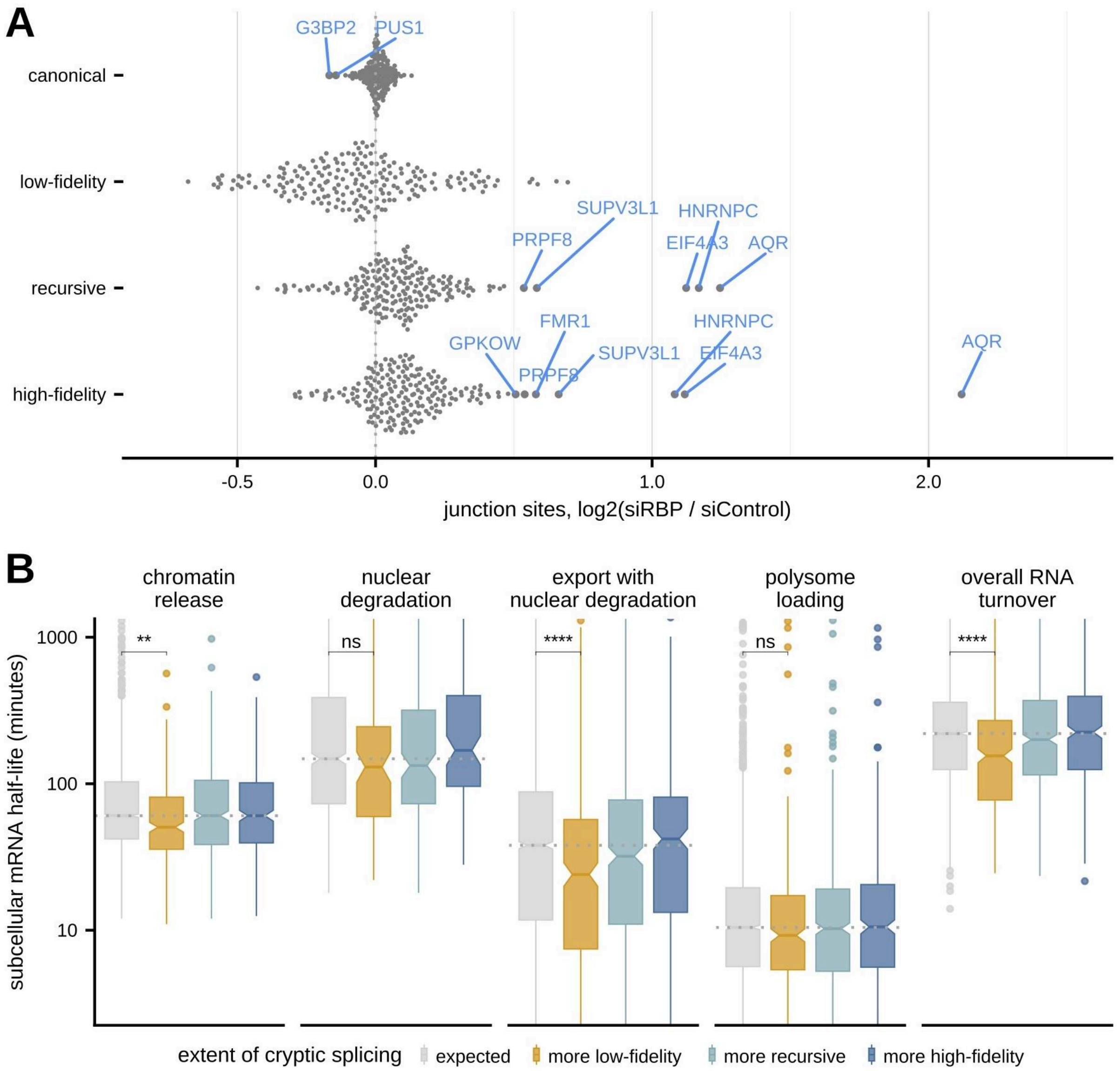


**Supplementary Figure 7. Identifying factors that influence cryptic splicing or the fate of cryptic transcripts. (A)** The fold change in junction sites between RNA binding protein siRNA knockdown and scrambled siRNA control samples (*x-axis*) for canonical, low-fidelity, recursive, and high-fidelity junction reads across all RBPs assessed by the ENCODE consortium. Significance was estimated using a permuted null distribution using control samples and significantly changing factors are labeled in blue (adjusted p-value < 0.05). **(B)** Distributions of mRNA residence times or half-lives across additional cellular compartments or fractions estimated using subcellular TimeLapse-seq from Ietswaart *et al.* 2024 for genes with more low-fidelity (*orange*), recursive (*teal*), or high-fidelity (*dark blue*) transcripts than predicted by the elastic net regression model, with genes whose cryptic splicing levels match expected levels shown in grey. Signficance was assessed with a Mann-Whitney U Test. **** adjusted p-value < 0.0001.

### Legends for Supplementary Tables

**Table S1. Sample-specific splice site identification.** Details about the number of reads, junction reads, and sites identified for each library across cell types and RNA enrichment methods.

**Table S2. Splice site-specific characterization.** Details about the position, quantification, sequence, conservation, motif fidelity, and pairing of canonical and cryptic splice sites identified across cell types and RNA enrichment methods.

**Table S3. Gene-specific splicing characterization.** Details about gene features, splice sites and reads, RNA abundance, and transcript abundance for canonical and cryptic splice sites identified across RNA enrichment methods in K562 cells.
